# Population-scalable genotyping from low-coverage sequencing data using pangenome graphs

**DOI:** 10.64898/2026.02.05.704023

**Authors:** Davide Bolognini, Andrea Guarracino, Chiara Paleni, Thomas S. Dudley, Licia Iacoviello, Alessandro Raveane, Peter H. Sudmant, Erik Garrison, Nicole Soranzo

## Abstract

Pangenome-based genotyping of structurally complex loci remains challenging at low sequencing depths, particularly when samples are of low quality, such as in ancient DNA. We present COSIGT (COsine SImilarity-based GenoTyper), a method that infers structural genotypes by matching short-read coverage patterns to pangenome haplotypes using cosine similarity. COSIGT maintains robust accuracy at low coverage (1-2X), outperforming existing methods where depth-sensitive approaches degrade. We demonstrate scalability to thousands of modern and ancient genomes, enabling population-scale analyses of complex variation from low-coverage data.

---

Structurally complex genomic regions, harboring multi-kilobase insertions, deletions, duplications, and copy-number variation, remain difficult to genotype accurately. A single linear reference cannot represent allelic diversity at these loci, and standard variant-calling workflows often fail to recover structural haplotypes [1]. Haplotype-resolved pangenomes address this limitation. The current human pangenome reference reveals thousands of previously inaccessible alleles at complex loci [2], improving read mapping and structural variant detection [3, 4].

Recent methods leverage pangenomes for targeted genotyping. Locityper aligns reads to locus-specific haplotypes and selects the best-fitting pair by optimizing alignment accuracy, insert-size concordance, and coverage balance [5]. This outperforms linear-reference methods at challenging loci. However, alignment-based likelihood models are sensitive to sequencing depth: as coverage decreases, signal-to-noise ratios degrade, reducing genotyping accuracy. This presents a barrier for low-coverage data, including population studies and biobank cohorts with variable depth [6]. Ancient DNA (aDNA) samples pose particular challenges, as coverage is often below 2X due to postmortem DNA degradation and microbial contamination [7].

Here, we describe COSIGT (COsine SImilarity-based GenoTyper), a pangenome-based pipeline for targeted genotyping at variable sequencing depths. The pipeline involves several steps (Figure 1A; Supplementary Figures S1–S3). COSIGT first constructs a variation graph of the target locus from resolved haplotypes (i.e., assemblies) with PGGB [8]. Building local graphs enables locus-specific parameter tuning, rapid incorporation of new assemblies, and faster genotyping compared to querying whole-genome pangenomes. Then, COSIGT maps sample’s reads to the local graph and determines which nodes they traverse, generating a read coverage vector over graph nodes. Each haplotype in the graph is represented as a coverage vector indicating traversal counts per node. To identify the most likely genotype, COSIGT enumerates all possible diploid haplotype pairs, constructs their expected coverage profiles by summing the corresponding haplotype vectors, and computes cosine similarities between each diploid profile and the observed sample vector. The synthetic haplotype pair yielding the highest similarity is selected as the predicted genotype. COSIGT’s implementation enables parallelization across genomic regions and samples, and provides visualizations to facilitate result interpretation (Figure 1B-D). Crucially, cosine similarity measures vector orientation rather than magnitude, making the metric inherently coverage-invariant and COSIGT less sensitive to sequencing depth than likelihood-based approaches.

**Fig. 1:**
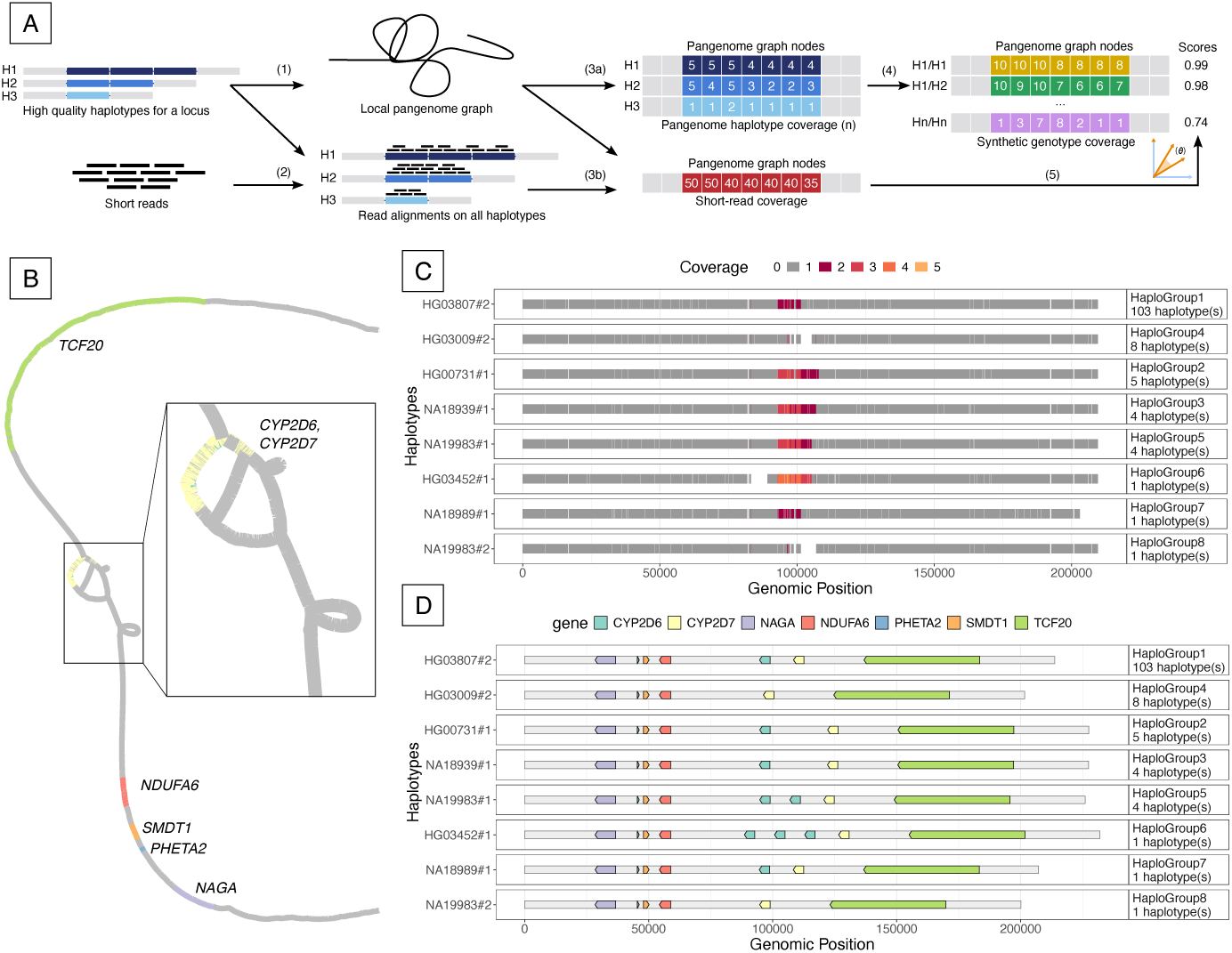
COSIGT genotyping workflow and CYP2D6/CYP2D7 locus example. A. Schematic of the COSIGT pipeline. 1) A local pangenome graph is built from input haplotypes. 2) Sequencing reads are aligned to local haplotypes. 3a) Haplotype coverage vectors are computed from graph node traversal counts. 3b) Read coverage vectors are computed from read-to-node alignments. 4) Synthetic diploid genotype vectors are generated for all possible haplotype pairs. 5) Cosine similarity between read and synthetic genotype vectors identifies the best-matching haplotype pair. **B. Local pangenome graph for the CYP2D6-CYP2D7 locus (chr22:42,031,864-42,245,566, GRCh38) visualized in Bandage [14].** Nodes are colored by gene content (legend in D). Multiple copies of CYP2D6 and CYP2D7 collapse onto shared nodes. **C. Node coverage heatmap for representative haplotypes at the CYP2D6-CYP2D7 locus.** Haplotypes are clustered by structural similarity, with one medoid per cluster shown. Variable coverage at CYP2D6/CYP2D7 nodes reflects distinct copy-number states across clusters. **D. Gene content for representative haplotypes.** Different clusters show distinct copy-number variation patterns.

We benchmarked COSIGT against Locityper on 326 challenging medically relevant genes (CMRGs) and 267 structurally variable regions (SVs) across multiple coverage levels in hundreds of short-read samples from 1000G consortium [9], using genome assemblies from the Human Pangenome Reference Consortium (HPRC) [2] and the Human Genome Structural Variation Consortium (HGSVC) [10] (Figure 2A). In leave-zero-out benchmarks, where true haplotypes are present in the graph, both methods achieve high accuracy at 5X and 30X, with *>* 93% high-quality calls. Locityper holds an advantage at 30X (98.2% vs. 93.9% in CMRGs; 98.2% vs. 96.7% in SVs). At low coverage, however, (1–2X), COSIGT substantially reduces error rates: 93.4% of calls reach mid-or-higher quality at 1X compared to 84.5% for Locityper, ; the advantage is maintained at 2X (95.8% vs. 93.4%) (Figure 2B,D; Extended Data Figure 1; Supplementary Figures S4–S7). In leave-all-out benchmarks, where true haplotypes are excluded from the pangenome, COSIGT remains near-optimal, achieving *>*= 87% of best-possible quality at most loci (87.6% of CMRGs, 90.8% of SVs) (Figure 2C; Extended Data Figure 2; Supplementary Figure S8). On simulated aDNA at 48 CMRGs enriched for pharmacogenetic and immunogenetic relevance, COSIGT consistently outperforms Locityper: at 1X it maintains ∼94% mid-or-higher quality while Locityper drops to ∼46%; at 2X COSIGT reaches ∼95% versus Locityper’s ∼60%. COSIGT attains higher accuracy than Locityper at all 48 genes (Figure 2D; Supplementary Figure S9).

**Fig. 2:**
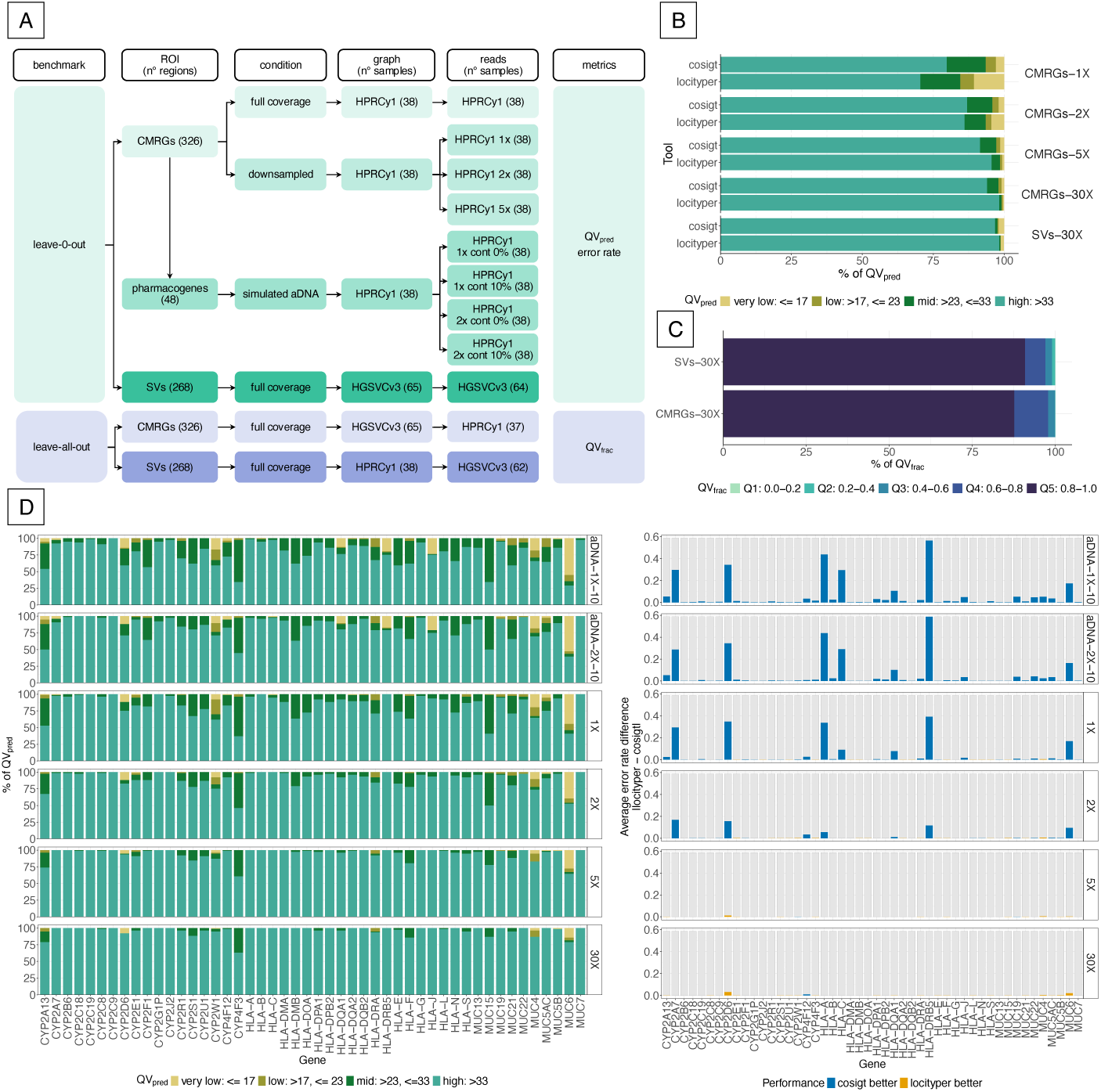
COSIGT benchmarking results. A. Benchmarking workflow. COSIGT in the leave-zero-out (ground-truth haplotypes present in graph) and leave-all-out (ground-truth haplotypes excluded) designs. Leave-zero-out: 326 CMRGs at 30X, 1X, 2X, and 5X coverage; 48 selected CMRGs with simulated ancient DNA (aDNA) (1X, 2X; 0% or 10% contamination); 265 SVs at 30X. Leave-all-out: CMRGs and 265 SVs at 30X. Pangenome graphs were built from haplotype-resolved assemblies (HPRC: 38 samples; HGSVC: 65 samples). Short-read samples matched graph samples (leave-zero-out) or were non-overlapping (leave-all-out). Performance metrics: alignment-based quality (*QV*_pred_) and error rate for leave-zero-out; fraction of maximum achievable quality (*QV*_frac_) for leave-all-out. **B. Leave-zero-out performance across coverage levels.** Stacked bar plots show *QV*_pred_ distribution for COSIGT and Locityper at CMRGs (1X, 2X, 5X, 30X) and SVs (30X). Quality categories: very low (*<* 17), low (17–23), mid (23–33), high (*>* 33). **C. Leave-all-out performance.** Stacked bar plots show *QV*_frac_ distribution for COSIGT at CMRGs and SVs (30X). Quintiles: Q1 (0.0–0.2), Q2 (0.2–0.4), Q3 (0.4–0.6), Q4 (0.6–0.8), Q5 (0.8–1.0). **D. Gene-level performance comparisons.** Left) *QV*_pred_ distribution for COSIGT on 48 selected CMRGs at 1X, 2X, 5X, and 30X coverage, plus simulated aDNA at 1X and 2X with 10% contamination. Right) Absolute error rate differences between methods across all genes, indicating which method performs better at each locus.

We further assessed COSIGT’s performance on 1,085 short-read wholegenome samples (∼20X coverage) from the Moli-sani cohort for HLA typing across several genes. COSIGT achieves a mean haplotype-level accuracy of 89.5% against HLA*IMP:02 imputed types [11], comparable to the dedicated HLA genotyping tool T1K (90.8%) [12] (Extended Data Figure 3; Supplementary Figure S10).

COSIGT addresses a key limitation of current genotyping methods: maintaining accuracy at low sequencing depth. Cosine similarity compares vector orientation independent of magnitude, making it robust to coverage variation. This is most valuable below 5X coverage, where COSIGT outperforms Locityper at most loci. At higher depths, both methods converge. Below 5X coverage, COSIGT outperforms Locityper at most loci, demonstrating particular utility for low-coverage aDNA data and large-scale biobanks with variable sequencing depth. The pipeline’s design is also compatible with long-read input, which could extend utility to low-coverage long-read datasets insufficient for assembly.

Some limitations merit consideration. Genotyping accuracy depends on the quality and completeness of the input pangenome: haplotypes absent from the reference panel cannot be called, and novel structural variants will be assigned to the most similar available haplotype. The leave-all-out benchmarks quantify this effect, showing that COSIGT still recovers a large fraction of maximum achievable accuracy when ground-truth haplotypes are absent from the pangenome. Additionally, COSIGT currently requires pre-aligned BAM/CRAM files, introducing a dependency on reference-based alignment. Future work will address this by enabling direct FASTQ input and implementing haplotype subsampling to maintain scalability as pangenome references grow.

Pangenome references are rapidly expanding in size, quality, and taxonomic breadth. As these resources grow, the limiting factor shifts from reference completeness to scalable genotyping. We have applied COSIGT to over 6,000 modern and ancient human genomes [13], demonstrating population-scale analysis of complex repeats and multi-copy genes (Supplementary Figures S11–S12). COSIGT’s tolerance for low coverage enables larger cohorts at reduced sequencing costs, inclusion of ancient samples, and analysis of datasets with heterogeneous depth, making the allelic diversity captured in pangenome references practically accessible for routine genotyping across the full spectrum of sequencing depths.

## Acknowledgments

The authors gratefully acknowledge support from Human Technopole and UC Berkeley IT teams for computational support. E.G. and A.G. are funded by NSF PPoSS Award #2118709; NIH R01HG013618; NIH U01HG013760; NIH U01DA057530; NIH U41HG010972; NIH R01HG013017; and the Tennessee Center for Integrative and Translational Genomics. The authors acknowledge Federica Santonastaso and Edoardo Giacopuzzi from Human Technopole for the initial processing of sample data of the Moli-sani cohort and support.

## Author Contributions Statement

E.G. conceived the project. D.B. and A.G. developed the software. D.B. wrote the documentation. D.B., A.G., C.P., and T.S.D. ran experiments. D.B. and C.P. made figures. A.R., L.I., P.H.S., E.G., and N.S. provided supervision. All authors wrote the manuscript.

## Methods

Here we provide details that are not described in the main text nor in other publications.

### Overview

COSIGT is a computational pipeline for structural genotype inference from sequencing data. The workflow is implemented in Snakemake v7 [16] to ensure reproducibility and scalability, and executes within isolated Singularity/Apptainer containers [17] or Conda environments [18], enabling consistent deployment across computing infrastructures.

COSIGT provides three workflow variants: the main pipeline for modern short-read samples (`master` branch), an `ancient dna` branch optimized for low-coverage and contaminated samples, and a `custom_alleles` branch for pre-extracted locus-specific sequences **(Supplementary Figures S1–S3)**. For the `master` and `ancient_dna` branches, the pipeline can be executed in two modes: refinement mode (`refine` subcommand) performs locus boundary optimization only, outputting refined BED coordinates for inspection before full genotyping; genotyping mode (`cosigt` subcommand) executes the complete workflow, using either user-provided or previously refined coordinates (see **Computational steps**). This two-stage approach enables iterative optimization of target regions for complex loci where initial boundary definitions may be suboptimal. The `custom_alleles` branch provides only the `cosigt` subcommand, as boundary refinement is not applicable when alleles are already extracted.

### Input Data

COSIGT requires four mandatory inputs:

1. *Chromosome-partitioned assemblies.* High-quality haplotype-resolved assemblies in FASTA format, partitioned by chromosome before processing **(Supplementary Data Note C.1)**. This reduces computational overhead by restricting analysis to relevant genomic regions. Alternatively, users can employ the `custom_alleles` branch of the pipeline, which accepts pre-extracted, locus-specific alleles in FASTA format for each target region, bypassing the need for whole-chromosome assemblies **(Supplementary Figure S2)**.
2. *Aligned short-read data.* Sequencing reads in BAM or CRAM format. COSIGT is compatible with large-scale public datasets such as the 1000 Genomes Project Phase 3 [9].
3. *Reference genome.* A reference genome sequence in FASTA format, used for (i) aligning input haplotypes to the corresponding reference chromosome and (ii) comparing structural alleles during genotyping. When CRAM files are provided, the reference genome is also required for read extraction.
4. *Target regions.* A BED file specifying genomic coordinates (reference-based) for loci to genotype. Optionally, columns 4 and 5 can specify locus names and alternative reference contigs mapping to the target regions, respectively (see **Short-read alignment and graph projection** and **Supplementary Data Note C.2**).

COSIGT accepts two optional inputs to improve genotyping quality and interpretability:

1. *Misassembly annotations.* A BED file specifying low-quality or misassembled regions in the input haplotypes (e.g., regions annotated as “Err”, “Dup”, or “Col” by the FLAGGER tool [2]). Haplotypes overlapping these regions are excluded from graph construction and genotyping to prevent artefacts.
2. *Gene annotations.* A GTF file with gene annotations and a FASTA file with translated protein sequences for the reference genome. When provided, COSIGT visualizes protein-coding gene content across selected haplotypes, enabling interpretation of structural variation at target loci.

All input files are automatically validated and organized by a dedicated Python script included with the pipeline, minimizing manual intervention (see the online documentation).

### Computational steps

#### Haplotype re-alignment

COSIGT initiates its workflow by aligning pre-partitioned, chromosome-specific haplotypes (query sequences) to their corresponding reference chromosome (target) using minimap2 [19] with the `asm20` preset, optimized for assembly-to-reference alignment. This preset terminates alignments when sequence divergence exceeds 20% but permits continuous alignment across less divergent segments within complex regions. This step precisely characterizes the relationship between sample-derived assemblies and the reference at target regions, which is essential for downstream extraction of structural haplotypes.

#### Region-of-interest extraction

Rather than constructing a full pangenome graph upfront, COSIGT applies an implicit pangenomics approach using impg (https://github.com/pangenome/impg), which treats wholegenome alignments generated by minimap2 as a queryable implicit graph.

##### Locus boundary refinement

Initial BED coordinates may not optimally capture structurally variable regions. The `refine` subcommand of impg adjusts boundaries to maximize haplotype coverage by iteratively expanding flanks (10 kb steps, up to 50% of locus length) and selecting coordinates where the most haplotypes span both edges. Alignments overlapping assembly-quality problems identified by FLAGGER are excluded. This refinement can be executed as a standalone preprocessing step via the refine subcommand of the pipeline, which outputs optimized coordinates for subsequent genotyping runs (see **Overview**).

##### Coordinate liftover and filtering

Target coordinates are projected onto all haplotypes using impg query, producing pairwise alignments in BEDPE format. These are filtered in three stages: (1) removal of alignments overlapping FLAGGER-flagged assembly regions; (2) merging of fragmented alignments within 200 kb to handle alignment gaps; and (3) retention of only those haplotypes whose merged alignment spans both locus boundaries (within 1 kb tolerance) as a single contiguous block. Haplotypes with multi-block alignments that may indicate misassembly or misalignment are excluded.

##### Sequence extraction

Filtered haplotype coordinates are used to extract sequences via bedtools [20], producing per-locus FASTA files for graph construction.

#### Region-of-interest pangenome graph construction

For each target region, PGGB [8] constructs a variation graph from the extracted haplotype sequences. PGGB performs all-to-all sequence alignment, unbiased graph induction [15], and progressive normalization to produce region-of-interest pangenome graphs in GFA format. The pipeline executes PGGB with default parameters except `--n-mappings 2` (to better support copy-number variable loci) and `--min-match-len 101` (to anchor graph construction on alignments of at least 101 bp). Graph statistics and manipulation are performed using ODGI [21]. The resulting pangenome graph merges all haplotypes into a unified structure where each haplotype corresponds to a path through the graph.

#### Recruitment of unmapped reads

To reduce reference bias, COSIGT employs kfilt (https://github.com/davidebolo1993/kfilt), a novel tool developed for this work that recruits sample reads failing to map to the reference genome but potentially aligning to alternative assemblies within each region-of-interest (ROI). The workflow proceeds as follows. First, Meryl [22] generates k-mer databases (`meryl count`, *k* = 31) for two sequence sets: the entire reference genome and the ROI comprising all haplotypes from the pangenome graph. From the ROI database, only k-mers unique to the ROI and absent from the rest of the reference genome are retained (`meryl difference`). Kfilt then builds a hybrid index from these ROI-specific k-mers using a tiered architecture that balances speed and sensitivity (`kfilt build -K 31`). A compact Bloom filter first provides *O*(1) rejection of k-mers absent from the database. K-mers passing this filter are queried against a hash table for *O*(1) exact and single-mismatch lookups. Finally, k-mers not found in the hash table are searched in a Burkhard-Keller (BK) tree for approximate matching at greater distances. This design minimizes costly tree searches, substantially improving throughput.

In practice, for each unmapped read, kfilt extracts all k-mers (*k* = 31) from both forward and reverse-complement sequences. Reads containing at least one exact k-mer match (Hamming distance = 0) are recruited for subsequent alignment (`kfilt filter -n 1 -m 0`). This stringent criterion ensures high specificity while recovering reads that share perfect k-mer matches with ROI haplotypes but diverge from the reference.

#### Short-read alignment and graph projection

Read-versus-graph alignment can be performed either by aligning reads directly to the graph or by mapping them to a linear pangenome reference and projecting these alignments onto the graph, as done in COSIGT. Short reads overlapping target regions are extracted from BAM/CRAM files with samtools [23], merged with unmapped reads recruited using kfilt (see **Recruitment of unmapped reads**), and aligned to haplotype sequences using bwa-mem2 [24]. To preserve informative multi-mapping, alignments are generated with an increased hit cap (-h 10000), retaining all hits whose score is at least 80% of the maximum for each read. These read-versus-sequence alignments are then projected onto the pangenome graph with gfainject (https://github.com/AndreaGuarracino/gfainject), which consumes the primary and alternative hits and reports graph-mapped alignments in GAF format.

When using reference genomes with alternate, unplaced, unlocalized, or patch scaffolds (such as the GRCh38 version used to align 1000 Genomes shortread data), an optional fifth column can be added to the target regions BED file to specify corresponding ALT contig coordinates for read recruitment. This ensures reads mapping to ALT scaffolds that represent the same genomic locus are captured alongside reads mapping to the primary chromosome assembly. COSIGT does not infer these extended regions automatically; the online documentation and **Supplementary Data Note C.2** provide instructions for generating such BED files during preprocessing.

#### Sample coverage vector computation

Coverage vector computation uses gafpack (https://github.com/pangenome/gafpack) to transform GAF alignments into per-node coverage features for each sample. For each graph node, gafpack computes the number of aligned bases normalized by node length (`--len-scale`), and further weights contributions from multi-mapping reads according to their number of occurrences (`--weight-queries`), yielding a coverage vector in which each element represents length-normalized, multi-mapping-aware coverage for that node in a given sample.

#### Haplotype coverage vector computation

Reference haplotype paths through the graph are converted into copy-number vectors (`odgi paths -H`), where each entry records how many times a given haplotype path traverses each graph node. For diploid samples, all possible unordered pairs of haplotype vectors are then summed to obtain candidate genotype vectors, which represent the expected node-wise copy number for each possible diploid haplotype combination (see **Genotype inference via cosine similarity**).

#### Haplotype clustering

Haplotypes are clustered into haplogroups using density-based spatial clustering (DBSCAN) [25]. Pairwise distances between haplotype paths are computed with odgi `similarity --distances` [21] and extracted from the `estimated.difference.rate` column, which quantifies dissimilarity as 1 − Dice, where Dice is the Sørensen–Dice coefficient derived from the Jaccard similarity of node coverage vectors. The resulting distance matrix is normalized by the maximum pairwise distance to scale all values between 0 and 1, and the mean normalized pairwise distance (mpd) is computed as a measure of overall region diversity.

An automated, data-driven procedure identifies the optimal *ε* parameter (maximum neighborhood radius for DBSCAN) by iteratively increasing it from 0.01 to 0.30 in steps of 0.01. Starting from *ε* = 0, where each haplotype forms its own cluster, the algorithm monitors cluster count stabilization: when the number of clusters changes by at most 1 between consecutive *ε* values, a plateau is detected, indicating that haplotypes have converged into natural groups. At this plateau, clustering stops if either of two conditions is met: (1) the number of clusters is at most *n*_haplotypes_*/*10, ensuring meaningful grouping rather than fragmentation into near-singletons; or (2) mpd exceeds 0.1, indicating a high-diversity region where natural haplogroups are expected to be more numerous. This adaptive stopping rule accommodates regional variation in haplotype diversity: in highly diverse regions, clustering terminates earlier to preserve distinct haplogroups, while in low-diversity regions, stricter consolidation is enforced to avoid over-splitting nearly identical haplotypes. If *ε* reaches 0.30 without meeting these criteria, it is capped at 0.30.

After clustering, the medoid of each haplogroup is identified as the haplotype with the smallest mean distance to all other members of that cluster. Haplogroups and their medoids are used in gene content and node coverage visualizations.

#### Node filtering

Nodes falling within low-complexity regions (for example, homopolymeric or repetitive sequences) or displaying outlier coverage values can degrade genotyping quality by introducing unreliable values into coverage vectors. These nodes are identified through a two-stage filtering process.

*Low-complexity nodes* are identified using Panplexity (https://github.com/AndreaGuarracino/panplexity). For each path in the graph, linguistic complexity is computed in 100 bp sliding windows (1 bp step) as the fraction of unique 16-mers relative to the theoretical maximum for that window. A global threshold is defined as *Q*_1_ − 1.5 × IQR, where *Q*_1_ is the first quartile and IQR is the interquartile range of all complexity scores across all paths (`panplexity -w 100 -d 100 -k 16 -t auto`). Any node overlapping a window below this threshold in any path is flagged.

*Coverage outlier nodes* are identified from haplotype path traversal counts. For each node, the mean copy number is computed across all haplotype paths, and nodes with values outside the range [*Q*_1_ − 1.5 × IQR*, Q*_3_ + 1.5 × IQR] are flagged as outliers.

Nodes flagged by either filter are excluded from downstream cosine similarity computation.

#### Genotype inference via cosine similarity

The core COSIGT algorithm computes cosine similarity between each sample’s read-derived coverage vector and all candidate diploid genotype vectors. For a sample coverage vector **s** (see **Sample coverage vector computation**) and a candidate genotype vector **g** (formed by summing two haplotype coverage vectors, see **Haplotype coverage vector computation**), cosine similarity is defined as:

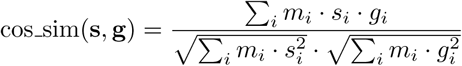

where *m_i_*∈ {0, 1} is a binary mask indicating whether node *i* passes the filtering criteria (see **Node filtering**): *m_i_*= 1 for nodes retained, *m_i_* = 0 for nodes flagged as low-complexity or coverage outliers.

This metric quantifies how closely the read coverage pattern matches each candidate diploid genotype while being scale-invariant to differences in read depth. The haplotype pair yielding the highest cosine similarity is selected as the predicted genotype. The method is implemented in Go (https://github.com/davidebolo1993/cosigt/blob/master/cosigt.go) and is computationally efficient, requiring only vector dot products and norms, and inherently tolerates sequencing noise and minor haplotype differences, as these perturbations reduce the similarity score gradually rather than preventing a genotype call entirely. While the current implementation supports diploid genotyping by enumerating all pairwise haplotype combinations, the framework generalizes to arbitrary ploidy *k* by enumerating *k*-tuples of haplotypes, though this extension is not yet integrated into the full pipeline.

### Alternate workflows

In addition to the `master` branch described so far **(Supplementary Figure S1)**, the COSIGT pipeline includes two alternative branches aimed at different use cases.

The `custom_alleles` branch allows the analysis to start from previously extracted haplotype sequences for each locus of interest, thus skipping the haplotype re-alignment and ROI extraction steps **(Supplementary Figure S2)**.

The `ancient_dna` branch replaces bwa-mem2 `mem` algorithm with bwa `aln` in the short-read alignment step, and skips the recruitment of unmapped reads step. Bwa-aln is run with parameters `-l 1024 -n 0.01 -o 2`, optimized for mapping ancient DNA [26] **(Supplementary Figure S3)**.

The `master` and `ancient_dna` branches can be run in `refine` mode to perform the locus boundary refinement analysis. This mode returns an adjusted list of loci with refined regions to maximize the number of retained haplotypes.

### Output Data

#### Main outputs

For each sample and locus, COSIGT reports a TSV file containing the sample identifier, predicted haplotype pair, haplogroup assignments, and the cosine similarity score of the best-matching genotype. All candidate haplotype pairs and their cosine similarity scores are saved in a separate file, ranked from highest to lowest similarity.

#### Visualization and quality control

COSIGT generates diagnostic visualizations to support structural variation analysis (1–3) and genotyping quality assessment (4–5). These include: (1) node coverage heatmaps for input haplotypes, grouped by haplogroup; (2) protein-coding gene content and orientation plots for input haplotypes, grouped by haplogroup; (3) pairwise alignments of predicted haplotypes to reference sequence; (4) read alignment coverage across all haplotypes; and (5) interactive plots comparing read and haplotype coverage vectors across graph nodes. Visualizations (1) and (2) are also generated for haplogroup medoids, with haplogroup frequencies annotated.

Node coverage heatmaps display graph structure and copy-number variation across haplotypes. Nodes are ordered by reference coordinates, and coverage values are capped at 12X. Coverage is rendered using segments, where the segment length corresponds to node length and color intensity reflects coverage depth, producing visualizations similar to `odgi viz` but with enhanced customization.

Gene content plots are generated when a gene annotation (GTF format) and reference protein sequences (FASTA format) are provided. Protein-coding sequences spanning the locus are aligned to all haplotypes using Miniprot [27]. Alignments are processed with Pangene [28], which constructs a gene graph accounting for copy-number variation, paralogs, and redundant annotations. The resulting BED output is visualized with the R package gggenes (https://github.com/wilkox/gggenes), displaying gene orientation and organization across haplotypes.

Haplotype alignment plots compare predicted haplotypes to reference sequences via all-versus-all minimap2 alignment. Alignments are visualized using the R package SVByEye [29] functions `plotAVA` and `plotMiro`, with aligned regions colored by percent identity in 1 kb bins.

Read alignment visualizations are generated using the `region` subcommand of Wally (https://github.com/tobiasrausch/wally). For each haplotype in the pangenome, all reads, regardless of their mapping quality (`-q 0`), are displayed across the full haplotype length with paired-end view enabled (`-p`), highlighting clipped reads (`-c`) and supplementary alignments (-u) to facilitate inspection of structural variant support and alignment quality.

Interactive coverage vector plots enable real-time comparison of read-derived and haplotype coverage patterns. A custom R Shiny application (see online documentation) accepts the COSIGT output folder as input and displays dual-axis scatter plots overlaying sample coverage (from read alignments) with candidate diploid genotype coverage (sum of two haplotype vectors) across graph nodes. Node positions are scaled by cumulative haplotype length, and users can interactively select haplotype pairs, toggle the filtering mask, and view cosine similarity scores and angular distances between vectors. This visualization aids in troubleshooting genotyping calls and assessing the contribution of individual nodes to the overall similarity metric.

### Benchmarking: Leave-zero-out and Leave-all-out

#### Assembly selection

Leave-zero-out and leave-all-out analyses utilized haplotype-resolved assemblies from two independent resources to ensure robustness across assembly methods and sample sets.

From the Human Genome Structural Variation Consortium phase 3 (HGSVCv3) [10], we downloaded 65 diploid assemblies (https://ftp.1000genomes.ebi.ac.uk/vol1/ftp/data_collections/HGSVC3) generated with Verkko [30]. One sample (HG00732) appeared in multiple batches; we retained only the most recent version from batch 3. Haplotype-phased FASTA files (haplotype 1, haplotype 2, and unassigned contigs) were converted to PanSN naming format (https://github.com/pangenome/PanSN-spec) with sample and haplotype identifiers to enable haplotype tracking in pangenome graphs.

From the Human Pangenome Reference Consortium year 1 (HPRCy1) [2], we used 38 diploid assemblies (maternal and paternal haplotypes) generated with hifiasm [31], selected from the updated year 2 release (HPRCy2; https://human-pangenomics.s3.amazonaws.com/index.html?prefix=submissions/DC27718F-5F38-43B0-9A78-270F395F13E8--INT_ASM_PRODUCTION) to match the original HPRCy1 sample composition. Chromosome naming was already PanSN-compliant and required no further processing. One original HPRCy1 sample (HG03492) is currently absent from the HPRCy2 release and was excluded.

Maternal and paternal haplotypes (and unassigned contigs for Verkko assemblies) were partitioned by chromosome using reference-based annotations provided by each consortium. For each chromosome partition in both assembly sets, the corresponding CHM13 reference chromosome (T2TCHM13v2.0; https://s3-us-west-2.amazonaws.com/human-pangenomics/T2T/CHM13/assemblies/analysis_set) was added to provide a complete telomere-to-telomere reference backbone for pangenome graph construction.

This dual-source design enabled assessment of COSIGT’s performance across assemblies generated with different long-read assembly algorithms (Verkko for HGSVCv3, hifiasm for HPRCy1). A complete list of downloaded assemblies with accession links and chromosome assignments is available in the paper repository (https://github.com/davidebolo1993/cosigt_paper/tree/main/resources/assemblies).

#### Short-read data

Matching short-read whole-genome sequencing data for benchmark samples were retrieved from the 1000 Genomes Project high-coverage release [9]. Sample identifiers and CRAM file locations were extracted from the available 1000 Genomes sequence index files (https://ftp.1000genomes.ebi.ac.uk/vol1/ftp/data_collections/1000G_2504_high_coverage/). All 38 HPRCy1 samples had matching highcoverage short-read data available in the 1000 Genomes collection. Among the 65 HGSVCv3 samples, 63 had matching data in the same repository. Sample NA24385, absent from the 1000 Genomes index, was retrieved from the European Nucleotide Archive (ENA) under project PRJEB35491. Sample NA21487 lacked publicly available short-read data and was excluded from benchmarking analyses. A complete list of downloaded alignment files with accession links is available in the paper repository (https://github.com/davidebolo1993/cosigt_paper/tree/main/resources/alignments).

#### Short-read realignment

1000 Genomes Project samples are aligned to a GRCh38 reference genome (https://ftp.1000genomes.ebi.ac.uk/vol1/ftp/technical/reference/GRCh38_reference_genome/) which is incompatible with Locityper, likely due to the presence of alternate contigs. Therefore, all short-read data were realigned to the GRCh38 primary assembly (GENCODE release 46: https://ftp.ebi.ac.uk/pub/databases/gencode/Gencode_human/release_46/). For each sample, the original CRAM or BAM file was name-sorted and converted to paired-end FASTQ files using samtools. Reads were then realigned to the GRCh38 primary assembly using bwa mem with parameters matching the original 1000 Genomes alignment protocol (-Y -K 100000000). Alignments were coordinate-sorted and compressed to CRAM format.

#### Leave-zero-out: CMRGs

We genotyped 38 short-read samples from the 1000 Genomes Project (see **Short-read data**) against their corresponding diploid assemblies from HPRCy1 (see **Assembly selection**), supplemented with CHM13 and GRCh38 reference assemblies as described above. A total of 326 target loci were derived from the Locityper benchmarking set [5], which extends the original list of 273 CMRGs [32] with additional genes of medical relevance, including complement factor H-related genes (CFHs), mucins (MUCs), cytochrome P450 genes (CYPs), and human leukocyte antigen genes (HLAs). Prior to genotyping, we performed locus boundary refinement analysis using the `refine` module of the COSIGT `master` branch and excluded haplotype blocks overlapping regions flagged by Flagger (https://github.com/human-pangenomics/hprc_intermediate_assembly/blob/main/data_tables/assembly_qc/flagger), which adjusted coordinates for 4 of these 326 loci (see **Locus boundary refinement**).

After genotyping with the `cosigt` module, all haplotypes were oriented to match the reference using `odgi flip`. Ground-truth haplotypes were identified by matching sample names to contig names in PanSN format. Genotyping accuracy was assessed using haplotype quality values (*QV*_pred_) as defined in the Locityper benchmarking framework:

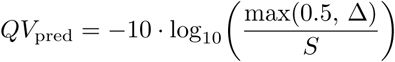

where Δ is the edit distance between predicted and expected haplotypes (with a minimum value of 0.5 to avoid undefined logarithms for perfect matches) and *S* is the alignment length, both computed using edlib [33].

The ratio Δ*/S* represents the error rate, which we used as a complementary metric to *QV*_pred_ for comparing methods. Because QV is logarithmic, equal QV differences represent different magnitudes of improvement depending on the quality range: for example, a 5-point QV increase from 10 to 15 reduces error rate from 10% to 3.2%, whereas the same 5-point increase from 30 to 35 reduces error rate from 0.1% to 0.032%, a 100-fold smaller absolute change. To facilitate direct comparison of method performance, we report both *QV*_pred_ distributions and absolute error rate differences (Figure 2D, Extended Data Figure 1; Supplementary Figures S4–S7).

For each sample, both possible pairings between predicted and expected haplotypes were evaluated, and the combination yielding the highest sum of QV scores was selected. QV metrics were computed only for samples with exactly two haplotypes retrieved during sequence extraction; sample counts per locus are reported in Extended Data Figure 1 and Supplementary Figures S4–S7. This evaluation approach is implemented in the `benchmark` module of the COSIGT `master` branch.

In addition to the full-coverage dataset (mean ∼30X, Supplementary Figure S6), short-read alignments were downsampled to mean coverages of 1X (Extended Data Figure 1), 2X (Supplementary Figure S4), and 5X (Supplementary Figure S5) using `samtools view` with `-s` subsampling fractions (`-s 42.0333, -s 42.0667, -s 42.1667`, respectively), and genotyping was repeated at each coverage level. A complete list of analyzed CMRG loci with per-sample genotyping results is available in the paper repository (https://github.com/davidebolo1993/cosigt_paper/tree/main/resources/benchmarking/leave_zero_out/cmrgs_predictions/modern).

#### Leave-zero-out: aDNA simulations of 48 CMRGs

We selected 48 CMRGs enriched for genes with pharmacogenetic and immunogenetic relevance (HLAs, CYPs, and MUCs) from the CMRG analysis. These gene families have been shaped by evolutionary pressures: HLAs exhibit long-standing balancing selection to maintain immune diversity [34], while CYPs and MUCs show signatures of local adaptation to environmental exposures and pathogens [35, 36].

For this set of loci, we simulated aDNA sequencing data from 38 HPRCy1 samples using ancestralSim (https://github.com/davidebolo1993/ancestralsim), a novel simulator developed for this study. AncestralSim uses Gargammel [37] to simulate ancient DNA sequencing reads from locus-specific pangenome haplotypes with realistic aDNA damage patterns. The pipeline extracts diploid haplotypes for each sample from the pangenome FASTA, simulates single-end reads with fragment length and deamination profiles characteristic of ancient DNA, trims adapters with fastp [38], and aligns reads to the corresponding reference chromosome using bwa aln with aDNA-optimized parameters (`-l 16500 -n 0.01 -o 2`). Exogenous DNA contamination is simulated by mixing reads from randomly selected non-self haplotypes at a specified ratio.

We simulated single-end reads at 1X and 2X coverage with mean fragment lengths of 70 bp (characteristic of degraded ancient DNA), read lengths of 100 bp, single-stranded deamination damage, and contamination levels of 0% or 10%. Since Locityper requires reads from a structurally simple background region on chromosome 17 for parameter estimation during preprocessing (see Locityper comparison), we additionally simulated diploid reads from the GRCh38 default region (chr17:72,062,001–76,562,000) for each sample. Region-specific BAM files were merged per sample to generate whole-genome alignments spanning the 48 target loci and the chromosome 17 background region.

Genotyping was performed using the `ancient_dna` branch of COSIGT, which substitutes bwa-mem2 `mem` algorithm with bwa aln and skips unmapped read recruitment (see **Alternate workflows**). Accuracy assessment followed the leave-zero-out protocol described for CMRGs, with *QV*_pred_ computed against ground-truth haplotypes from HPRCy1 assemblies (Supplementary Figure S9). Genotyping results are available in the paper repository (https://github.com/davidebolo1993/cosigt_paper/tree/main/resources/benchmarking/leave_zero_out/cmrgs_predictions/aDNA).

#### Leave-zero-out: SVs

We genotyped 64 short-read samples from the 1000 Genomes Project (see **Short-read data**) against 65 diploid assemblies from HGSVCv3 (see **Assembly selection**), supplemented with CHM13 and GRCh38 reference assemblies. Flagger-flagged regions (https://ftp.1000genomes.ebi.ac.uk/vol1/ftp/data_collections/HGSVC3/working/20241218_phase3-main-pub_data/uwash/flagger/verkko/final_beds_alt_removed/) were excluded as described for CMRGs. Target loci were defined by identifying protein-coding genes overlapping structural variants larger than 10 kb in the HPRCy1 pangenome. Each region was extended by 100 kb on both sides using bedtools `slop`, and overlapping intervals were merged with bedtools `merge`. We excluded regions located on sex chromosomes and those longer than 500 kb, with the exception of the glycophorin locus (GYP), which was retained despite exceeding the length threshold. These regions were processed with locus boundary refinement analysis, after which we discarded FBLN2, BTNL3, and MYOM2 due to insufficient haplotype coverage (fewer than half the maximum possible number of haplotypes fully spanning those regions). These filtering steps yielded a final set of 265 loci. Genotyping and accuracy assessment followed the same protocol described for CMRGs, with *QV*_pred_ calculated for samples with exactly two retrieved haplotypes. The analysis was performed at 30X coverage (Supplementary Figure S7). A complete list of analyzed SV loci with per-sample genotyping results is available in the paper repository (https://github.com/davidebolo1993/cosigt_paper/tree/main/resources/benchmarking/leave_zero_out/svs_predictions).

#### Leave-all-out: CMRGs

To assess COSIGT’s performance when groundtruth haplotypes are absent from the reference panel, we performed a leave-all-out analysis in which short-read samples were genotyped using pangenome graphs built exclusively from non-overlapping assembly sets. The HPRCy1 and HGSVCv3 sample sets share two individuals; these were removed from the short-read data for leave-all-out analysis. We genotyped the remaining 36 HPRCy1 short-read samples against pangenome graphs built from 65 HGSVCv3 assemblies (supplemented with CHM13 and GRCh38). The 326 CMRG loci from the leave-zero-out analysis (see **Leave-zero-out: CMRGs**) were used as target regions.

For samples with exactly two ground-truth haplotypes retrieved in the corresponding leave-zero-out analysis, we computed: (1) *QV*_pred_, the observed quality obtained by comparing COSIGT-predicted haplotypes to the ground-truth haplotypes (as defined in the leave-zero-out analysis); (2) *QV*_max_, the maximum achievable quality obtained by comparing each ground-truth haplotype to all haplotypes in the leave-all-out graph and selecting the best match; and (3) *QV*_frac_ = *QV*_pred_*/QV*_max_, quantifying how closely COSIGT approaches the best possible accuracy given the haplotype diversity available in the graph. *QV*_max_ represents the theoretical upper bound on genotyping accuracy given the available reference panel. For each ground-truth haplotype, we exhaustively computed QV scores against all haplotypes in the graph and selected the highest-scoring match. The sum of the two best-matching QV scores defines *QV*_max_ for that sample. A *QV*_frac_ of 1.0 indicates that COSIGT selected the most similar available haplotypes; lower values indicate suboptimal selections relative to the graph content. For visualization, we binned *QV*_frac_ values into quintiles (Q1: 0.0–0.2, Q2: 0.2–0.4, Q3: 0.4–0.6, Q4: 0.6– 0.8, Q5: 0.8–1.0) and plotted the percentage of loci falling within each quintile (Figure 2C and Extended Data Figure 2). Per-sample genotyping results are available in the paper repository (https://github.com/davidebolo1993/cosigt_paper/tree/main/resources/benchmarking/leave_all_out/cmrgs_predictions).

#### Leave-all-out: SVs

We genotyped 62 HGSVCv3 short-read samples (excluding the two samples overlapping with HPRCy1) against pangenome graphs built from 38 HPRCy1 assemblies, supplemented with CHM13 and GRCh38. The 265 SV loci from the leave-zero-out analysis were used as target regions. *QV*_pred_, *QV*_max_, and *QV*_frac_ were computed as described for the leave-all-out CMRG analysis (Supplementary Figure S8). Per-sample genotyping results are available in the paper repository (https://github.com/davidebolo1993/cosigt_paper/tree/main/resources/benchmarking/leave_all_out/svs_predictions).

#### Runtime and memory usage

We evaluated the computational requirements of the COSIGT pipeline using Snakemake’s built-in benchmarking functionality on the leave-zero-out analysis of 265 SV loci (see **Leave-zero-out: SVs**). This generated *>*126,000 individual benchmark measurements across pipeline rules executed on multiple samples, chromosomes, and genomic regions. Benchmark data were aggregated per rule to compute performance statistics (minimum, mean, median, and maximum) for execution time and memory consumption (Supplementary Figures S11–S12).

The analysis revealed substantial heterogeneity in computational demands across pipeline stages. Pangenome graph construction with PGGB exhibited the widest performance range (median: 5.01 min, range: 3.28–70.64 min; median memory: 1,257 MB, range: 668–7,758 MB). Sample-specific genotyping steps demonstrated consistently low resource requirements: cosigt tool (median: 0.020 min, range: 0.0097–0.35 min; median memory: 52 MB, range: 0.7–176 MB), gfainject (median: 0.55 min, range: 0.29–14.73 min), and gafpack (median: 0.19 min, range: 0.11–4.69 min). Assembly-to-reference alignment with minimap2 showed the greatest variability (median: 3.77 min, range: 0.81–70.76 min; median memory: 9,396 MB, range: 2,031–28,787 MB), reflecting differences in assembly contiguity and chromosome size. Reference k-mer database construction with meryl required 17.8 GB memory but represents a one-time preprocessing cost amortized across all samples.

To estimate the theoretical minimum runtime, we performed critical path analysis by constructing a directed acyclic graph of rule dependencies and computing the longest path assuming unlimited parallelization. Under this idealized scenario, the critical path would require approximately 168.6 minutes (∼2.8 hours), dominated by assembly alignment (70.8 min) and pangenome construction (70.6 min). This theoretical minimum represents the time required when all parallelizable steps execute simultaneously and does not account for practical constraints such as available CPU cores, memory limits, or job scheduling overhead. This analysis demonstrates that the pipeline’s architecture enables substantial parallelization across genomic regions and samples, making it well-suited for large-scale population genomic analyses.

Computational requirements of the COSIGT pipeline are summarized in the paper repository (https://github.com/davidebolo1993/cosigt_paper/tree/main/plot/data/time_mem_benchmark).

#### Locityper comparison

We benchmarked COSIGT against Locityper v1.2.0 [5] using the same CMRG and SV loci and short-read samples described above. For each analysis, we extracted haplotype sequences from the COSIGT pipeline output and provided them to Locityper as the reference haplotype database. Locityper was run following the standard workflow described in the documentation (https://locityper.vercel.app/commands): database construction (`locityper add`), sample preprocessing (`locityper preproc`), and genotyping (`locityper genotype`). For aDNA simulations, the chromosome 17 background region required for Locityper preprocessing was included in the simulated BAM files as described above. Additionally, for aDNA datasets, the preproc command was run with `--tech illumina` to override automatic read length detection, which misidentified the sequencing platform due to the short fragment sizes characteristic of aDNA.

*QV*_pred_ values for Locityper were computed by comparing predicted haplotypes (extracted from the resulting JSON files) to ground-truth haplotypes using the same edlib-based approach described for COSIGT. Haplotype sequences were extracted from the reference-oriented pangenome graph generated by `odgi flip` to ensure consistent coordinate systems. For each sample, both possible pairings between predicted and expected haplotypes were evaluated, and the combination yielding the highest sum of QV scores was selected. Locityper genotyping was performed at coverage levels of 1X, 2X, 5X, and 30X for CMRGs, and at 30X only for SVs. For aDNA simulations, Locityper was run at 1X and 2X coverage with 0% and 10% contamination. Per-sample genotyping predictions for Locityper are provided side-by-side with COSIGT predictions in the corresponding paper repository folders.

### Benchmarking: HLA typing

#### Sample selection and alignment

We selected 1,085 samples for HLA typing from 6,676 high-quality short-read samples in the Moli-sani cohort. The complete cohort consisted of 6,811 samples sequenced on an Illumina NovaSeq 6000 (150PE mode, mean 20X coverage) using Illumina’s PCR-free library kit. Sequencing was performed in four batches: 4,428 samples by an external provider and three batches (2,068, 282, and 33 samples) by the Human Technopole National Facilities. The first batch was pre-processed and aligned to GRCh38.p12 using dragen v05.121.645.4.0.3 (Illumina). Remaining batches were pre-processed with fastp [38] and aligned to iGenomes 2023.1 GATK GRCh38 using bwa. All samples underwent duplicate marking with GATK MarkDuplicates [39] v4.3.0.0 and variant calling with DeepVariant (WGS model, implemented in NVIDIA Parabricks) [40, 41].

#### Quality control

After computing QC metrics (mosdepth [42], samtools, somalier [43], SCE-VCF v0.1.2 [44], bcftools [45]), samples were excluded based on the following quality control criteria: less than 80% of bases with ≥10X coverage; mean coverage on any autosome *<*15X; CHARR (contamination from homozygous alternate reference reads) *>*0.03, or *>*15% heterozygous genotypes with inconsistent allele balance; mismatch between WGS-inferred sex and metadata; failure to match microarray genotyping data (bcftools gtcheck); duplicate samples (relatedness ≥0.9, excluding known twin pairs); Ts/Tv ratio *<*1.9 or *>*2.1. These criteria excluded 135 samples from the original 6,811, yielding 6,676 high-quality samples.

#### Pangenome construction

HLA typing analyses used 234 assemblies from HPRCy2 (https://human-pangenomics.s3.amazonaws.com/index.html?prefix=submissions/DC27718F-5F38-43B0-9A78-270F395F13E8--INT_ASM_PRODUCTION), supplemented with GRCh38. Target regions for seven classical HLA genes (HLA-A, -B, -C, -DPA1, -DPB1, -DQA1, -DQB1) were defined and extended to include corresponding alternate reference contigs, capturing reads mapping to ALT scaffolds.

#### HLA type annotation

Haplotype sequences extracted by COSIGT were annotated with ImmuAnnot using the reference database from Zenodo (https://zenodo.org/records/10948964). ImmuAnnot assigns HLA types based on exon-level sequence similarity between predicted haplotypes and known alleles. The two predicted haplotype types were combined to produce the genotype for each Moli-sani sample.

#### T1K comparison

We compared COSIGT with T1K v1.0.8-r237 [12] using identical alignment files and HLA-annotated haplotype sequences as input. T1K genotyping followed the standard workflow with parameters: `--preset hla-wgs --alleleDelimiter : --skipPostAnalysis.`

#### Microarray-based validation

Array genotyping was performed using a custom Affymetrix SNP array IDEA-900k, designed in collaboration with Thermo Fisher. Among the selected markers, 10,600 were specifically added across HLA loci. COSIGT and T1K predictions were validated against 4-digit HLA typing performed with HLA*IMP:02 [11] on the array genotypes. For each HLA gene, we computed sample-level accuracy (both alleles correct) and haplotype-level accuracy (individual allele correct). Samples were excluded from comparison if they: (1) failed HLA*IMP:02 imputation, (2) failed T1K genotyping, or (3) carried HLA types absent from the HPRCy2 reference panel (Extended Data Figure 3). We additionally recomputed sample- and haplotype-level accuracies after removing samples with T1K genotyping quality ≤ 0, following the authors’ recommendations (https://github.com/mourisl/T1K) (Supplementary Figure S10). HLA typing comparison statistics are available in the paper repository (https://github.com/davidebolo1993/cosigt_paper/tree/main/resources/benchmarking/hla_typing).

## Data Availability

COSIGT-predicted genotypes can be found at https://doi.org/10.5281/zenodo.18477735. Sample-level and raw data for the Moli-sani cohort are available under controlled access.

## Code Availability

COSIGT is available at https://github.com/davidebolo1993/cosigt. COSIGT and Locityper predictions for all datasets analyzed in this study, along with scripts to reproduce the statistics and figures, are available at https://github.com/davidebolo1993/cosigt_paper. Documentation is available at https://davidebolo1993.github.io/cosigtdoc.

## Reporting summary

Further information on research design is available in the Nature Portfolio Reporting Summary linked to this article.

## Supplementary Material

### Supplementary A Extended Data Figures

**Fig. 1:**
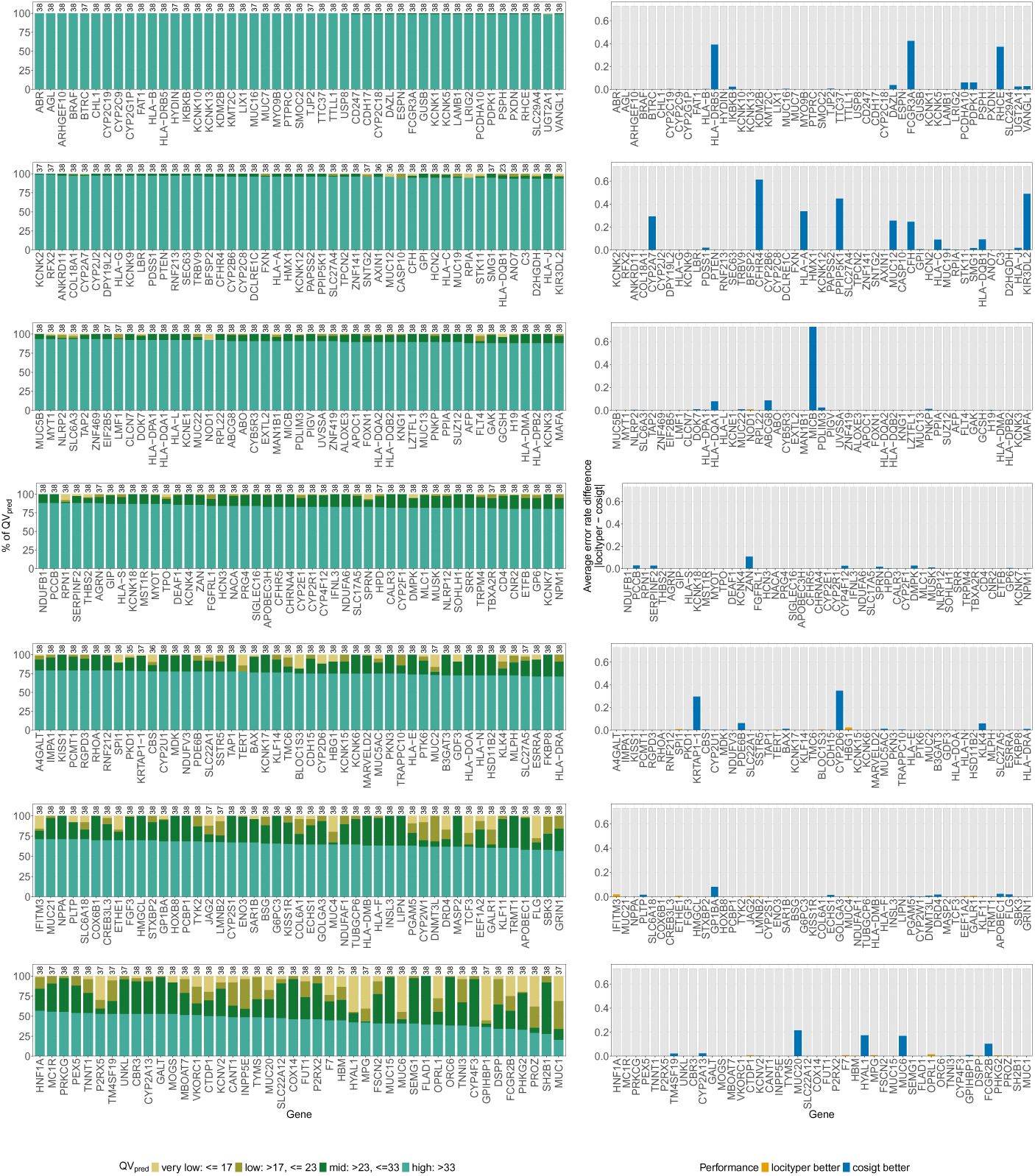
Gene-level COSIGT performance in leave-zero-out analysis of CMRGs at 1X coverage. Per-gene comparison of COSIGT and Locityper genotyping accuracy for 326 CMRGs at 1X coverage. Left: *QV*_pred_ distribution for COSIGT, showing the percentage of samples in each quality category (very low: <17, low: 17–23, mid: 23–33, high: >33; see Methods). Sample counts per gene are indicated above each bar. Right: Absolute error rate differences between COSIGT and Locityper. Genes are ordered by COSIGT performance (higher to lower percentage of high QV calls per gene).

**Fig. 2:**
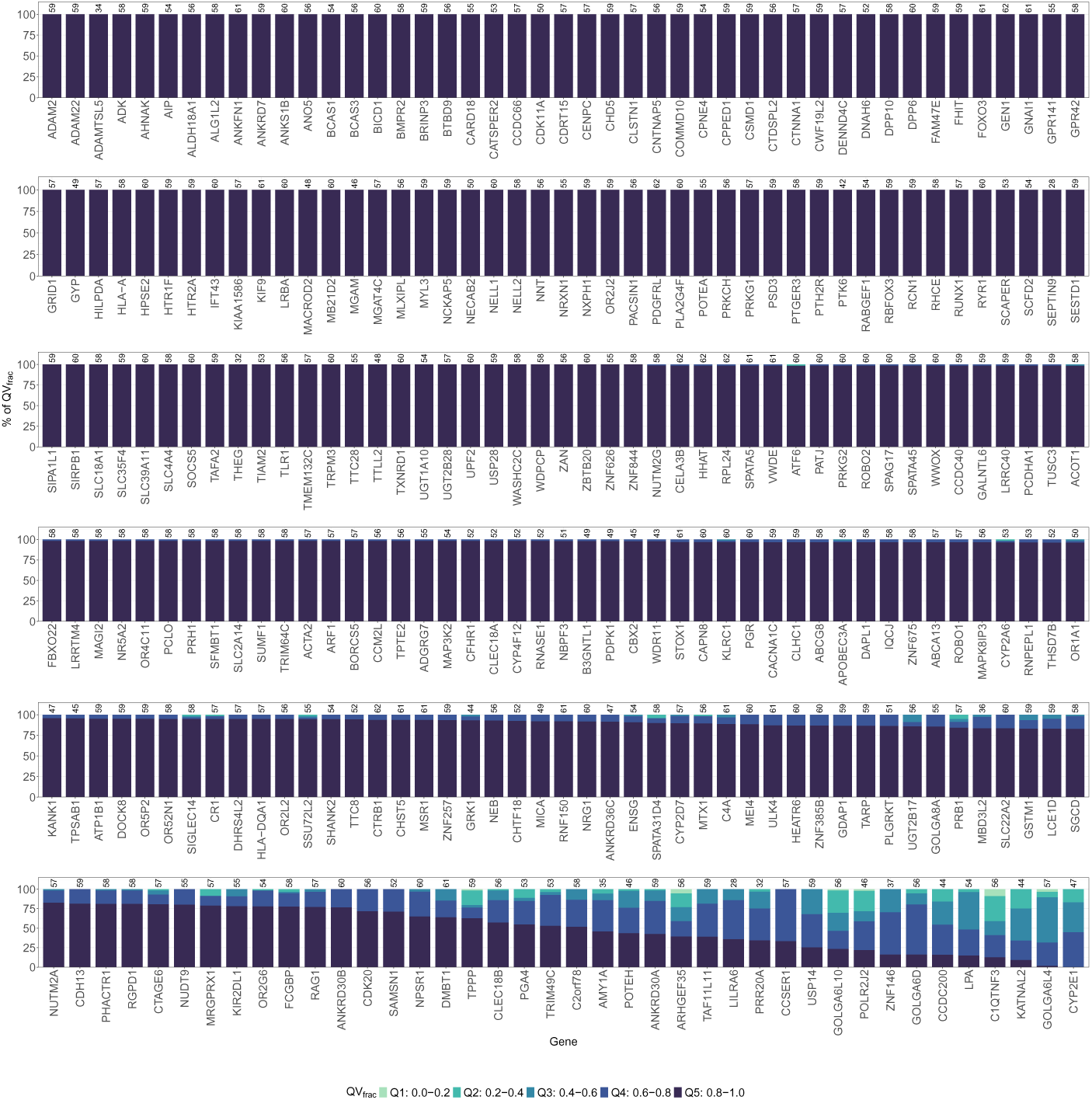
Gene-level COSIGT performance in leave-all-out analysis of SVs at 30X coverage. Per-gene distribution of *QV*_frac_ for 265 SVs in the leave-all-out setting. *QV*_frac_ quantifies how closely COSIGT approaches the best possible accuracy given the available haplotype diversity in the reference panel, with values binned into quintiles (Q1: 0.0–0.2, Q2: 0.2–0.4, Q3: 0.4–0.6, Q4: 0.6–0.8, Q5: 0.8–1.0; see Methods). Genes are ordered by COSIGT performance (higher to lower percentage of Q5 calls per gene). Sample counts per gene are indicated above each bar.

**Fig. 3:**
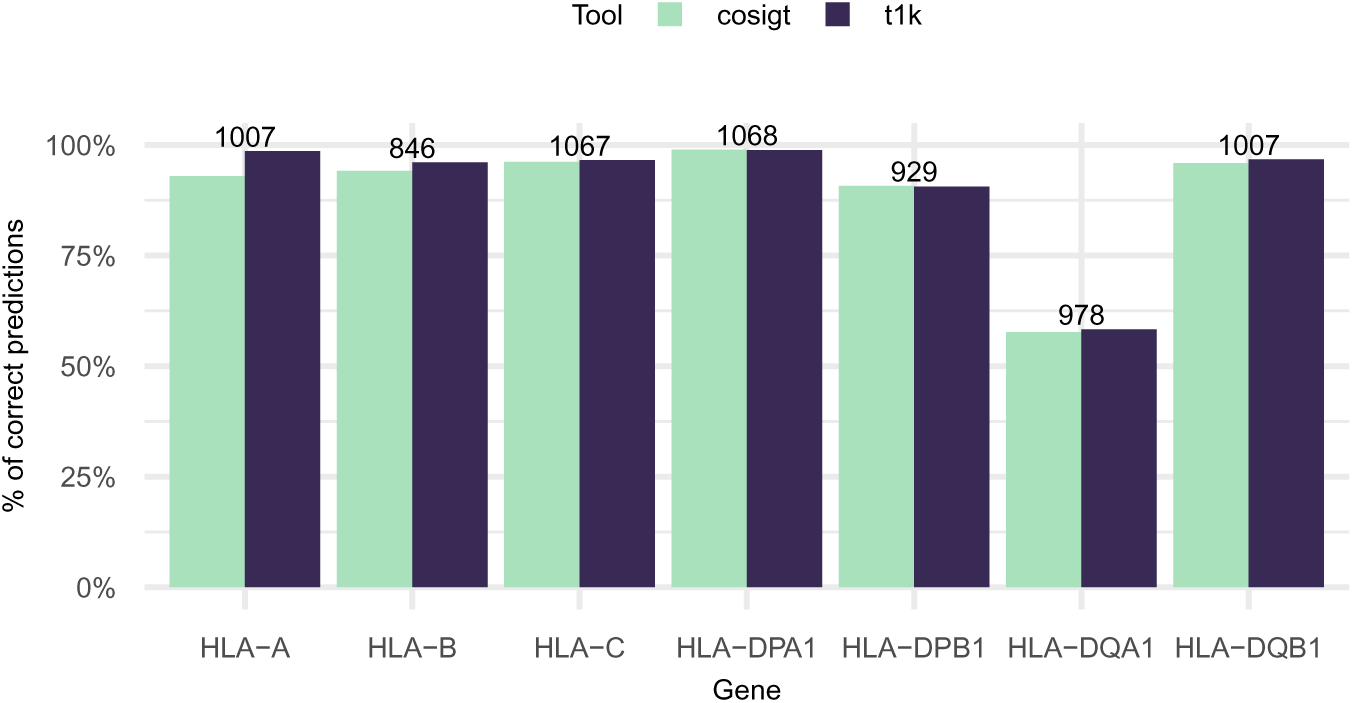
COSIGT performance on HLA typing. Haplotype-level concordance between short-read-based genotyping (COSIGT, T1K) and microarray-based imputation (HLA*IMP:02) for seven HLA genes (HLA-A, -B, -C, -DPA1, -DPB1, -DQA1, -DQB1) in a subset (*n* = 1,085) of the Moli-sani cohort. Numbers above bars indicate samples with predictions from all three methods and at least one reference haplotype present in the pangenome graph for the imputed genotype. See Methods for details.

### Supplementary B Supplementary Figures

**Fig. S1:**
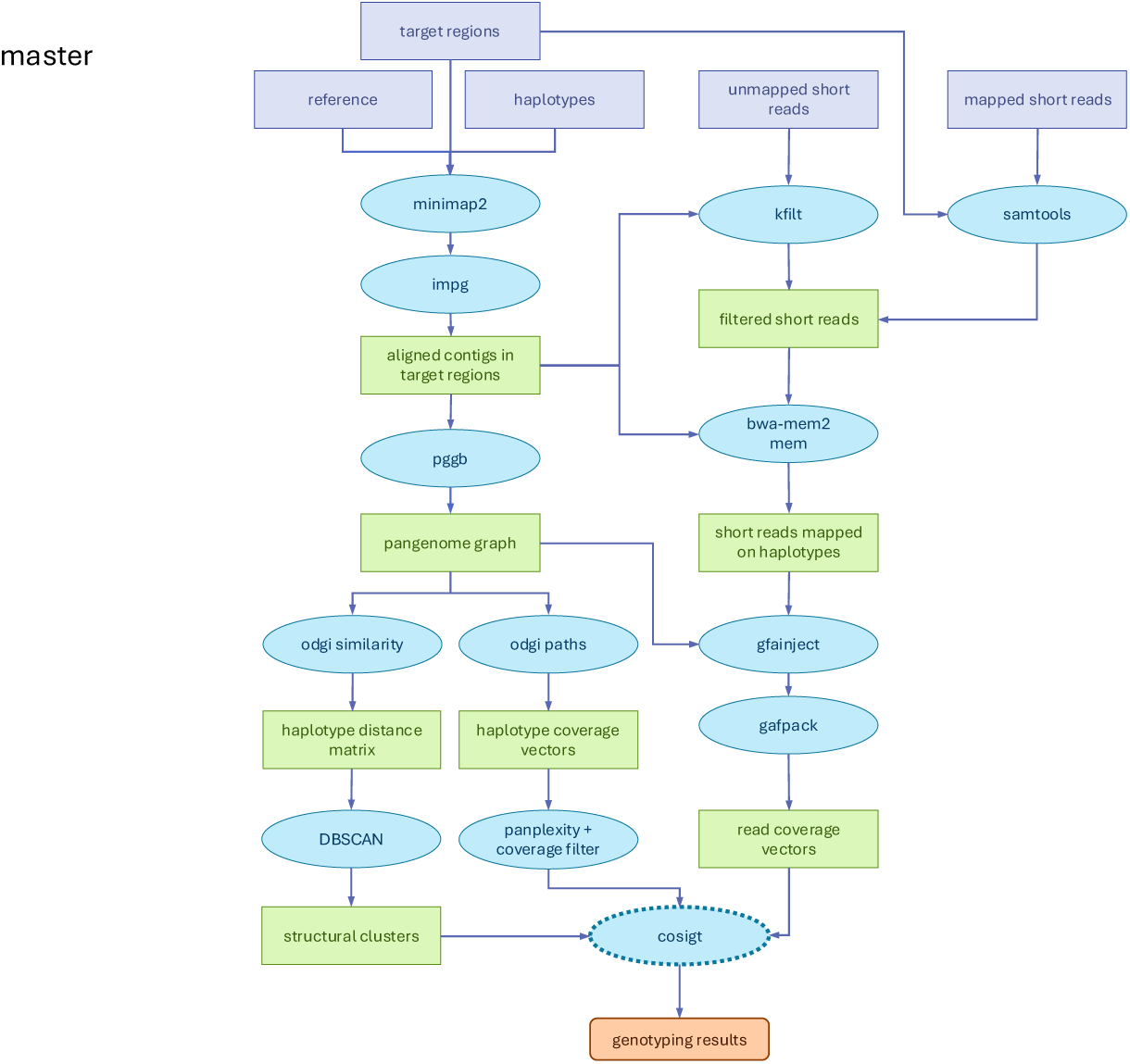
DAG of the COSIGT master pipeline branch. Simplified directed acyclic graph (DAG) describing the `master` branch of the COSIGT pipeline, for the cosigt subcommand. Only required inputs and main outputs are represented. Purple rectangles: required input files. Blue ovals: main tools utilized by the pipeline. Green rectangles: main intermediate results. Orange rounded rectangle: final genotyping results. See Methods.

**Fig. S2:**
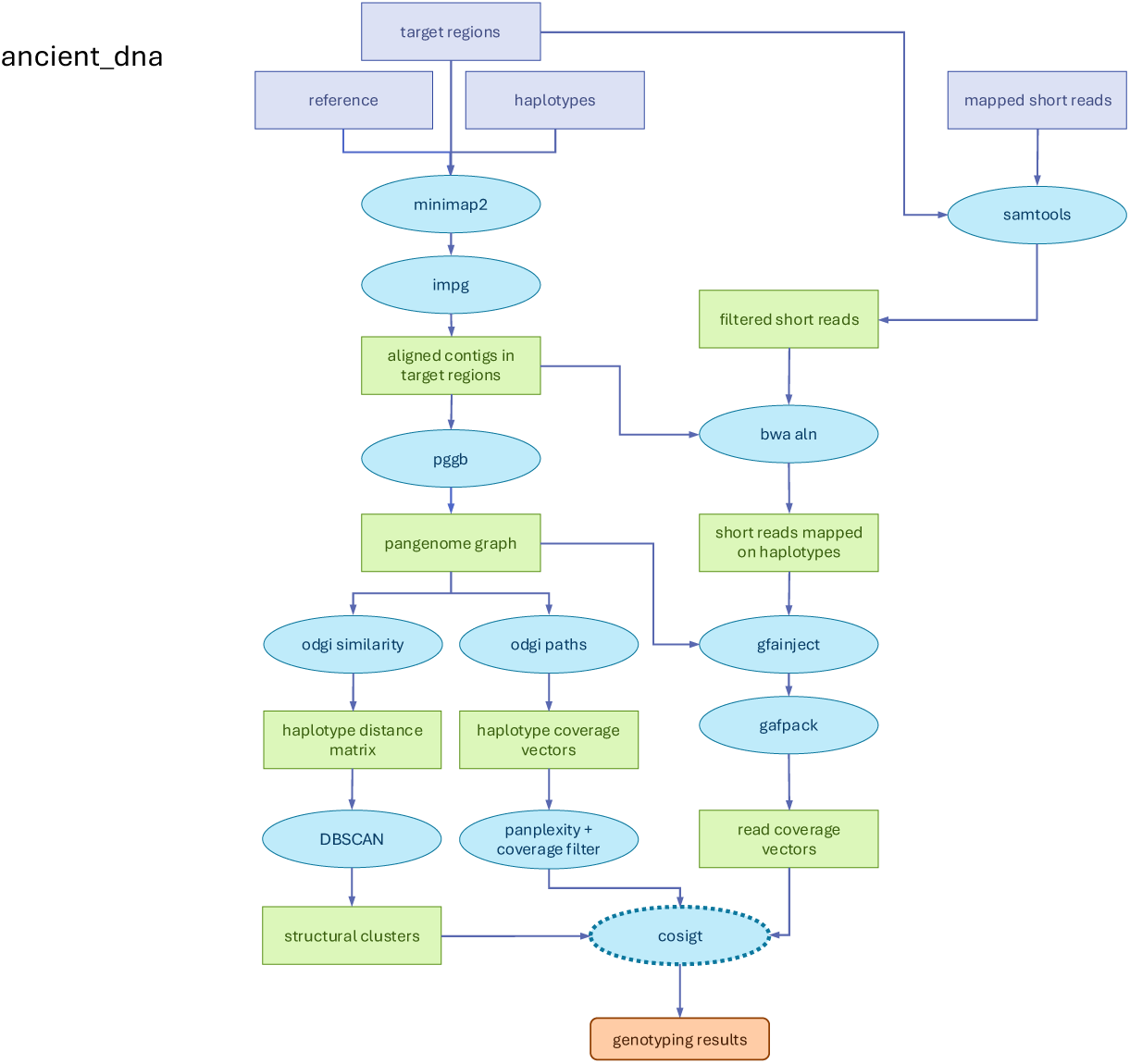
DAG of the COSIGT custom_alleles pipeline branch. Simplified DAG describing the `custom_alleles` branch of the COSIGT pipeline for the cosigt subcommand. Compared to the `master` branch, `custom_alleles` skips the alignment of contigs to the target regions and exploits user-provided alleles for graph building and read alignment. See Methods.

**Fig. S3:**
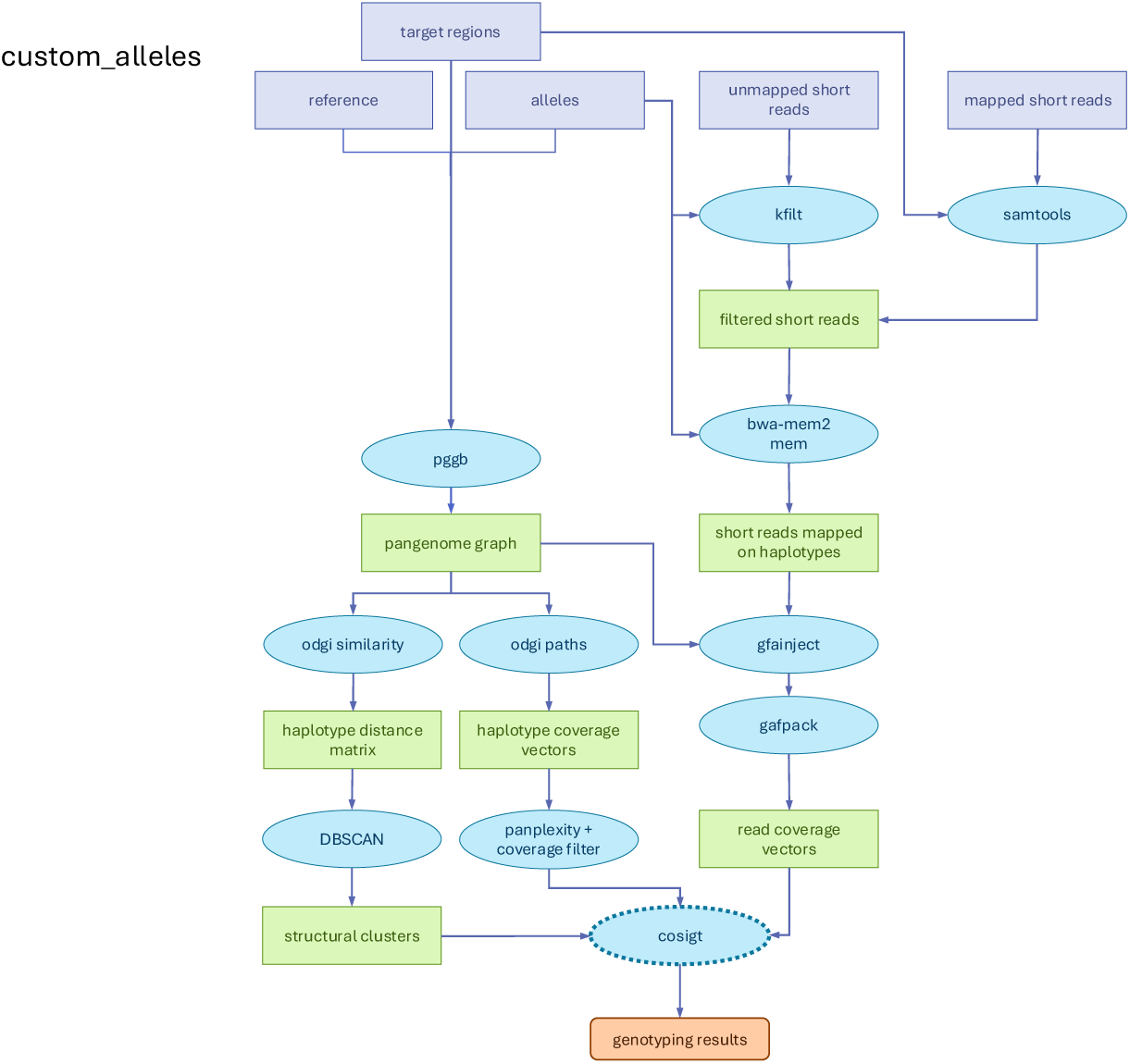
DAG of the COSIGT ancient_dna pipeline branch. Simplified DAG describing the `ancient_dna` branch of the COSIGT pipeline for the cosigt subcommand. Compared to the `master` branch, this branch skips the recruitment of unmapped reads and replaces bwa-mem2 `mem` algorithm with bwa `aln` for read-to-haplotype alignment. See Methods.

**Fig. S4:**
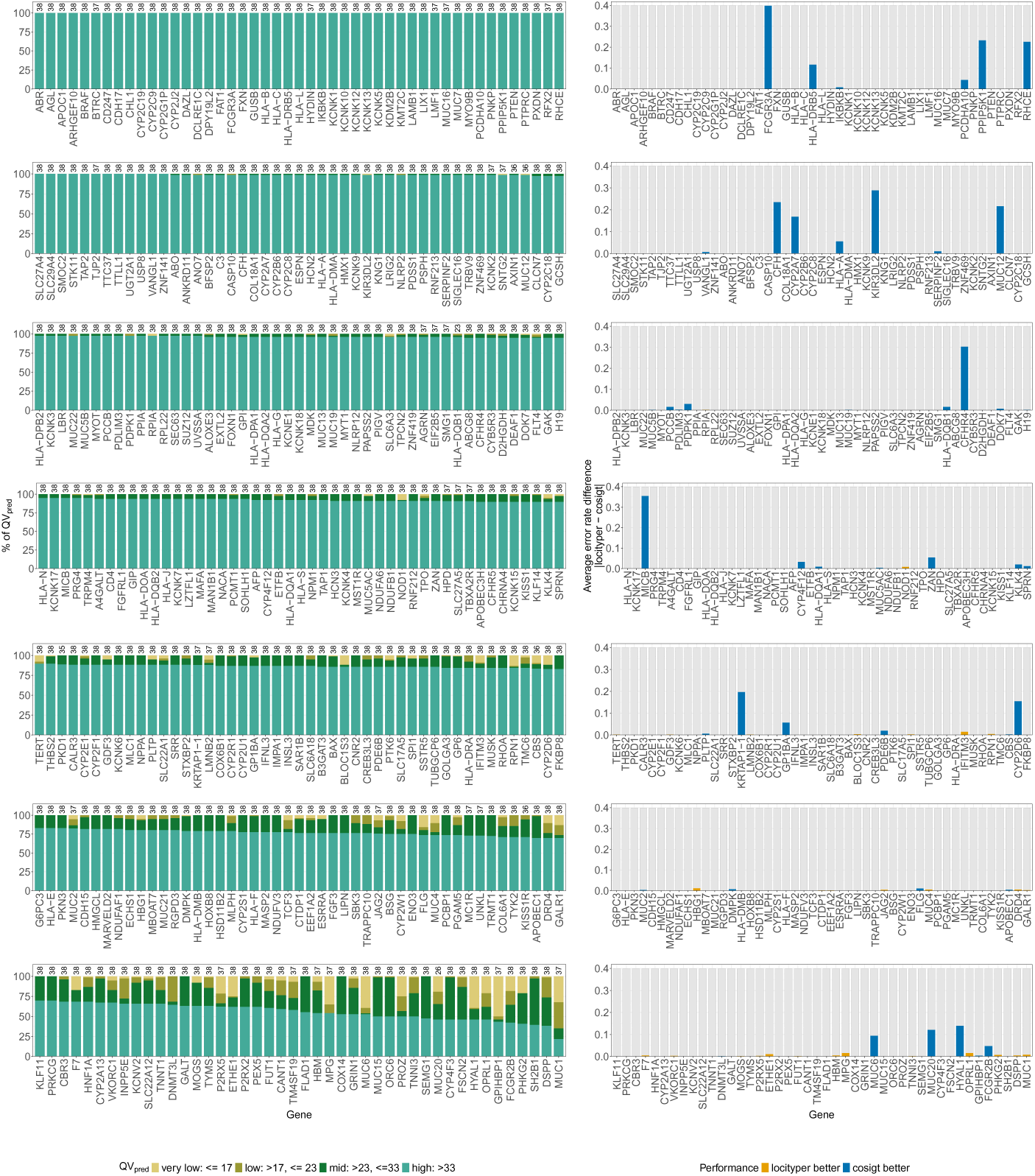
Gene-level COSIGT performance in leave-zero-out analysis of CMRGs at 2X coverage. Per-gene comparison of COSIGT and Locityper genotyping accuracy for 326 CMRGs at 2X coverage. Left: *QV*_pred_ distribution for COSIGT, showing the percentage of samples in each quality category (very low: <17, low: 17– 23, mid: 23–33, high: >33; see Methods). Sample counts per gene are indicated above each bar. Right: Average absolute error rate differences between COSIGT and Locityper. Genes are ordered by COSIGT performance (higher to lower percentage of high QV calls per gene).

**Fig. S5:**
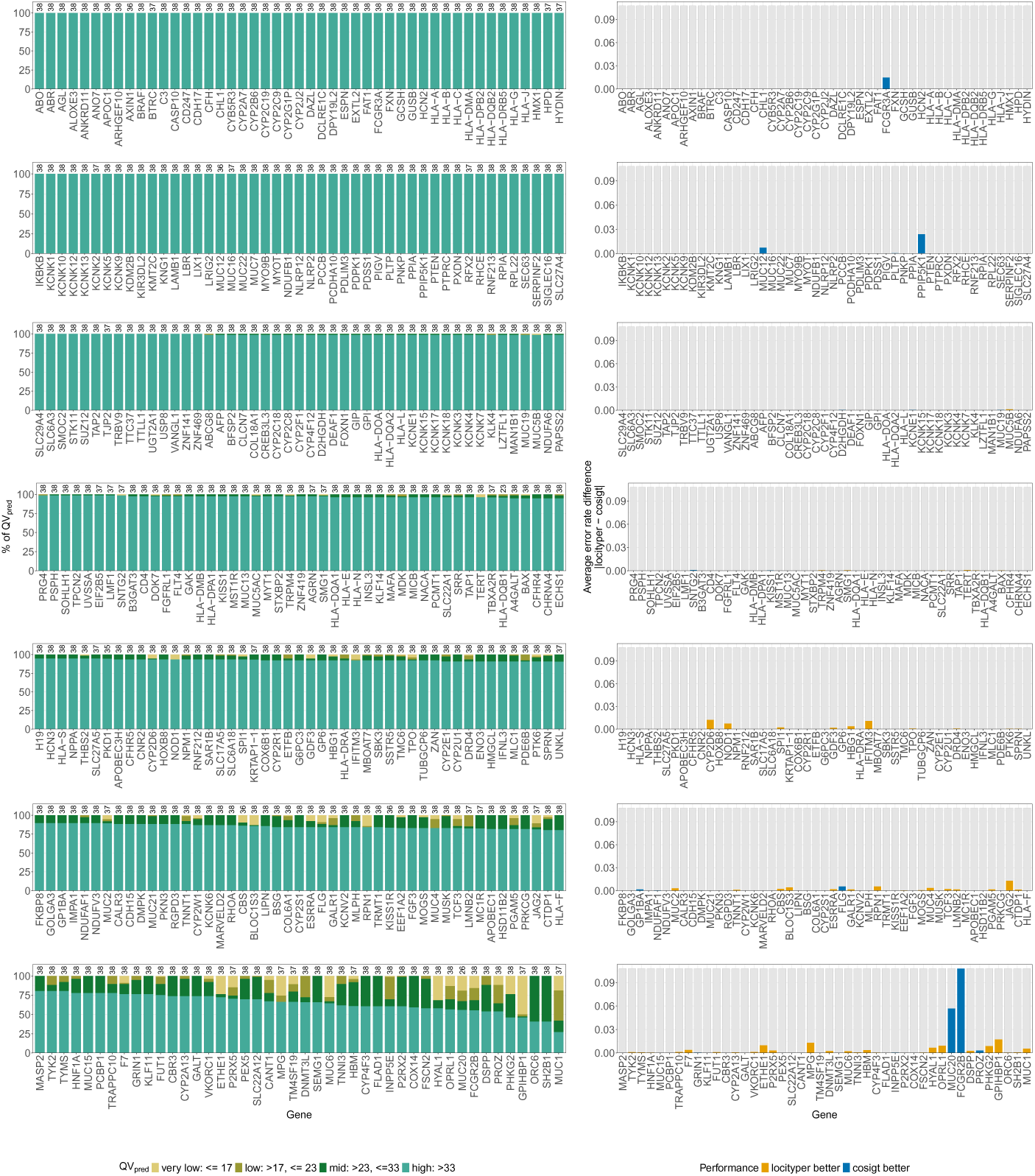
Gene-level COSIGT performance in leave-zero-out analysis of CMRGs at 5X coverage. Per-gene comparison of COSIGT and Locityper genotyping accuracy for 326 CMRGs at 5X coverage. Left: *QV*_pred_ distribution for COSIGT, showing the percentage of samples in each quality category (very low: <17, low: 17– 23, mid: 23–33, high: >33; see Methods). Sample counts per gene are indicated above each bar. Right: Average absolute error rate differences between COSIGT and Locityper. Genes are ordered by COSIGT performance (higher to lower percentage of high QV calls per gene).

**Fig. S6:**
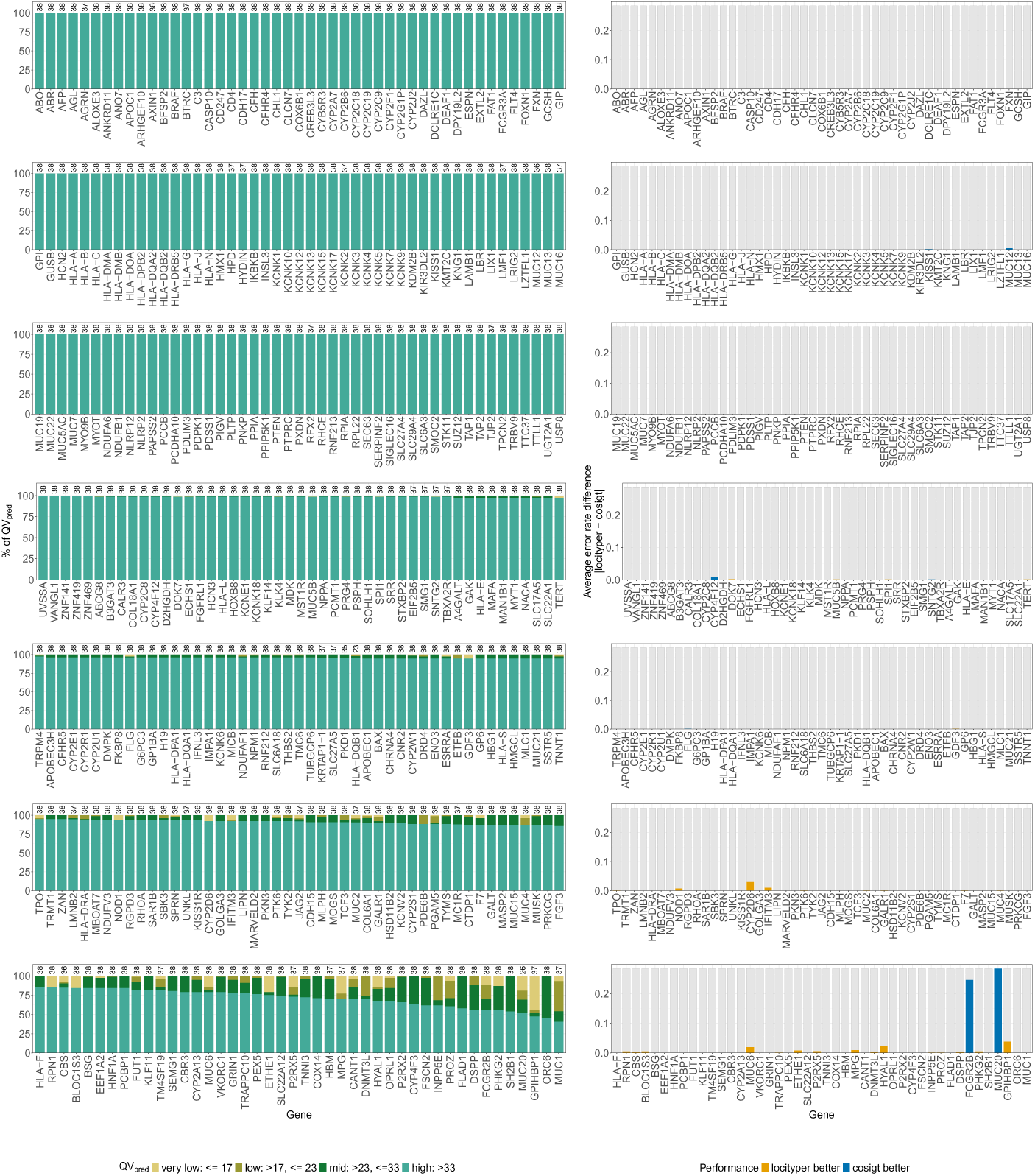
Gene-level COSIGT performance in leave-zero-out analysis of CMRGs at 30X coverage. Per-gene comparison of COSIGT and Locityper genotyping accuracy for 326 CMRGs at 30X coverage. Left: *QV*_pred_ distribution for COSIGT, showing the percentage of samples in each quality category (very low: <17, low: 17–23, mid: 23–33, high: >33; see Methods). Sample counts per gene are indicated above each bar. Right: Average absolute error rate differences between COSIGT and Locityper. Genes are ordered by COSIGT performance (higher to lower percentage of high QV calls per gene).

**Fig. S7:**
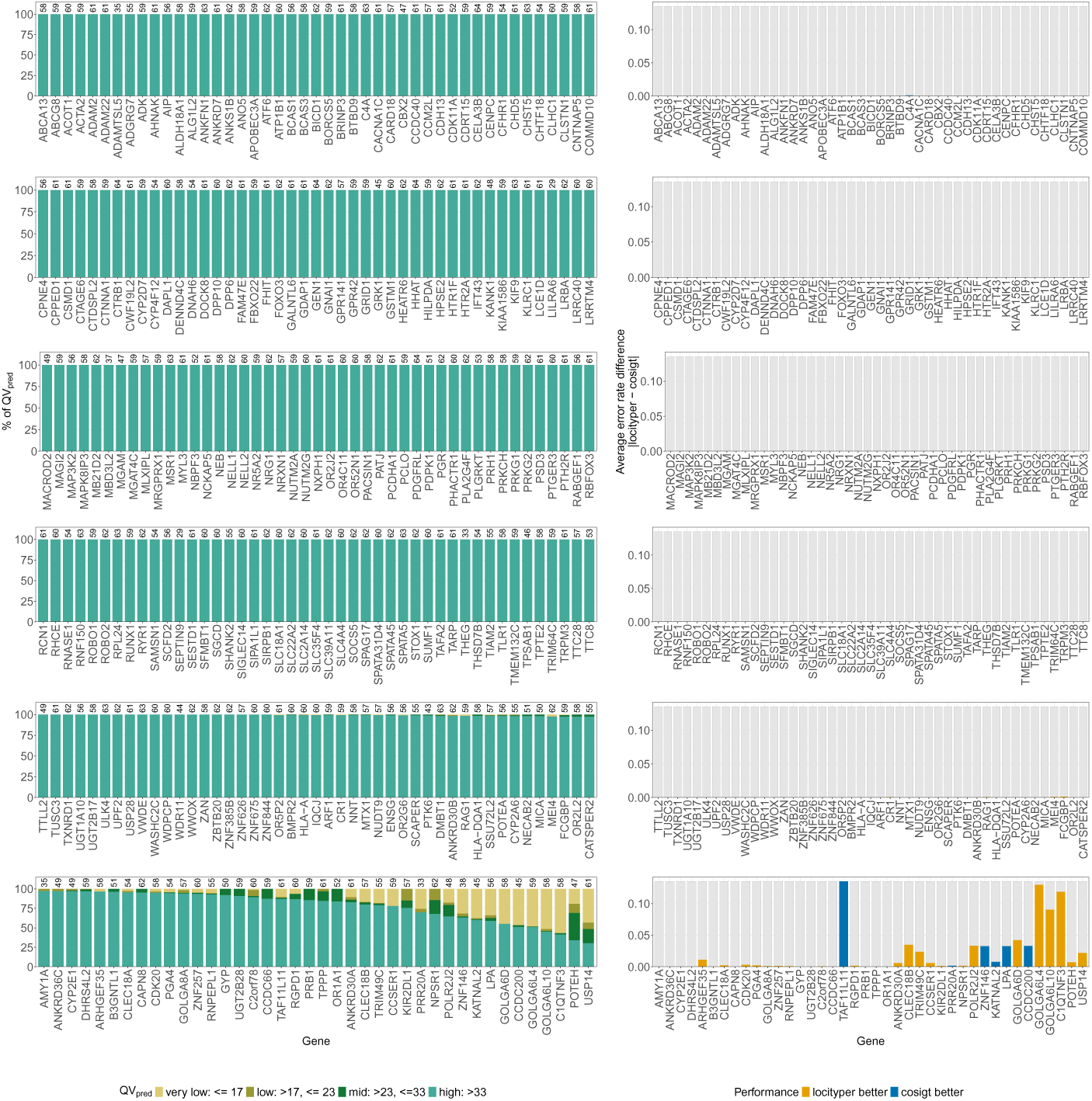
Gene-level COSIGT performance in leave-zero-out analysis of SVs at 30X coverage. Per-gene comparison of COSIGT and Locityper genotyping accuracy for 265 SVs at 30X coverage. Left: *QV*_pred_ distribution for COSIGT, showing the percentage of samples in each quality category (very low: <17, low: 17–23, mid: 23–33, high: >33; see Methods). Sample counts per gene are indicated above each bar. Right: Average absolute error rate differences between COSIGT and Locityper. Genes are ordered by COSIGT performance (higher to lower percentage of high QV calls per gene).

**Fig. S8:**
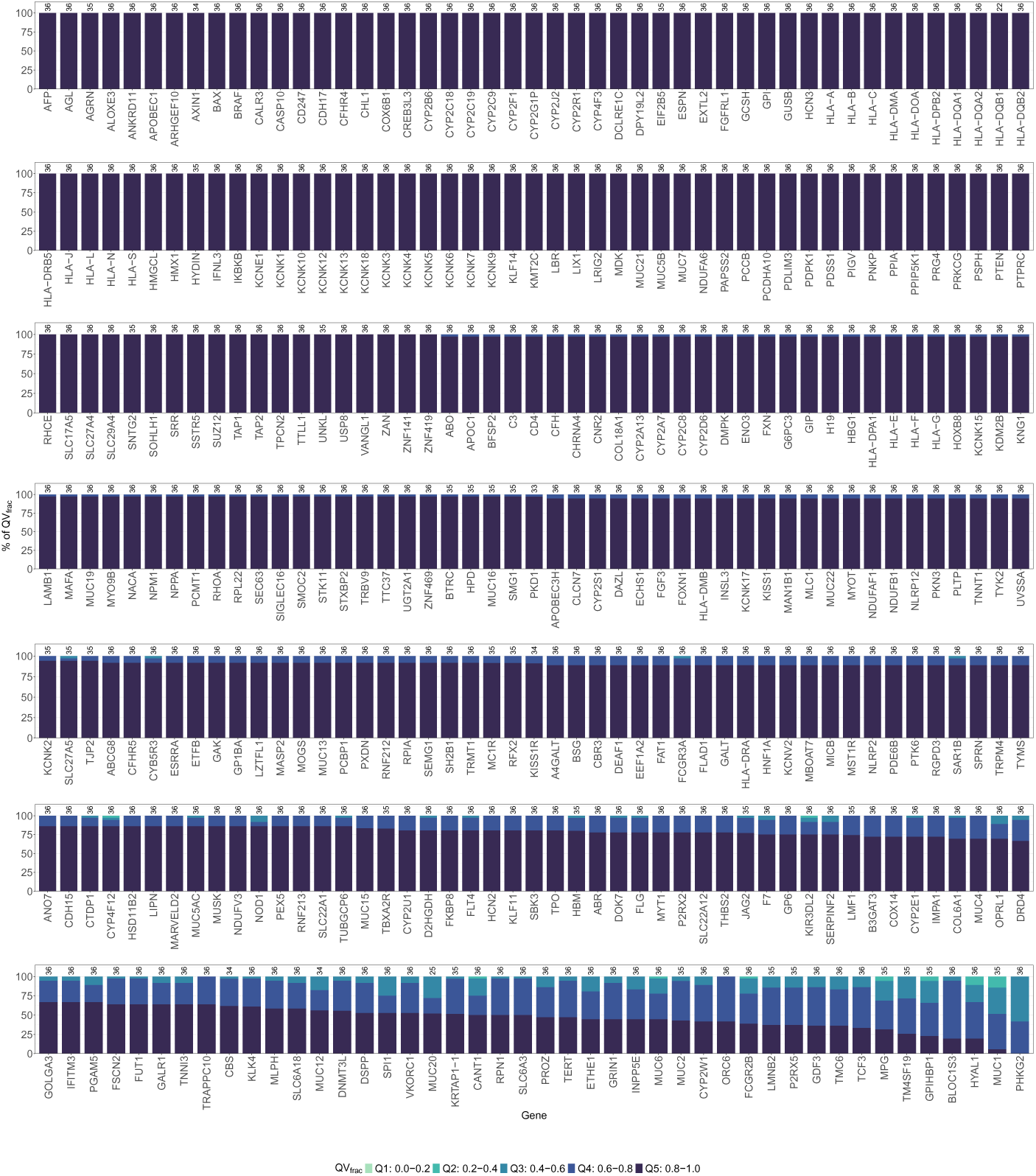
Gene-level COSIGT performance in leave-all-out analysis of CMRGs at 30X coverage. Per-gene distribution of *QV*_frac_ for 326 CMRGs in the leave-all-out setting. *QV*_frac_ quantifies how closely COSIGT approaches the best possible accuracy given the available haplotype diversity in the reference panel, with values binned into quintiles (Q1: 0.0–0.2, Q2: 0.2–0.4, Q3: 0.4–0.6, Q4: 0.6–0.8, Q5: 0.8–1.0; see Methods). Genes are ordered by COSIGT performance (higher to lower percentage of Q5 calls per locus). Sample counts per gene are indicated above each bar.

**Fig. S9:**
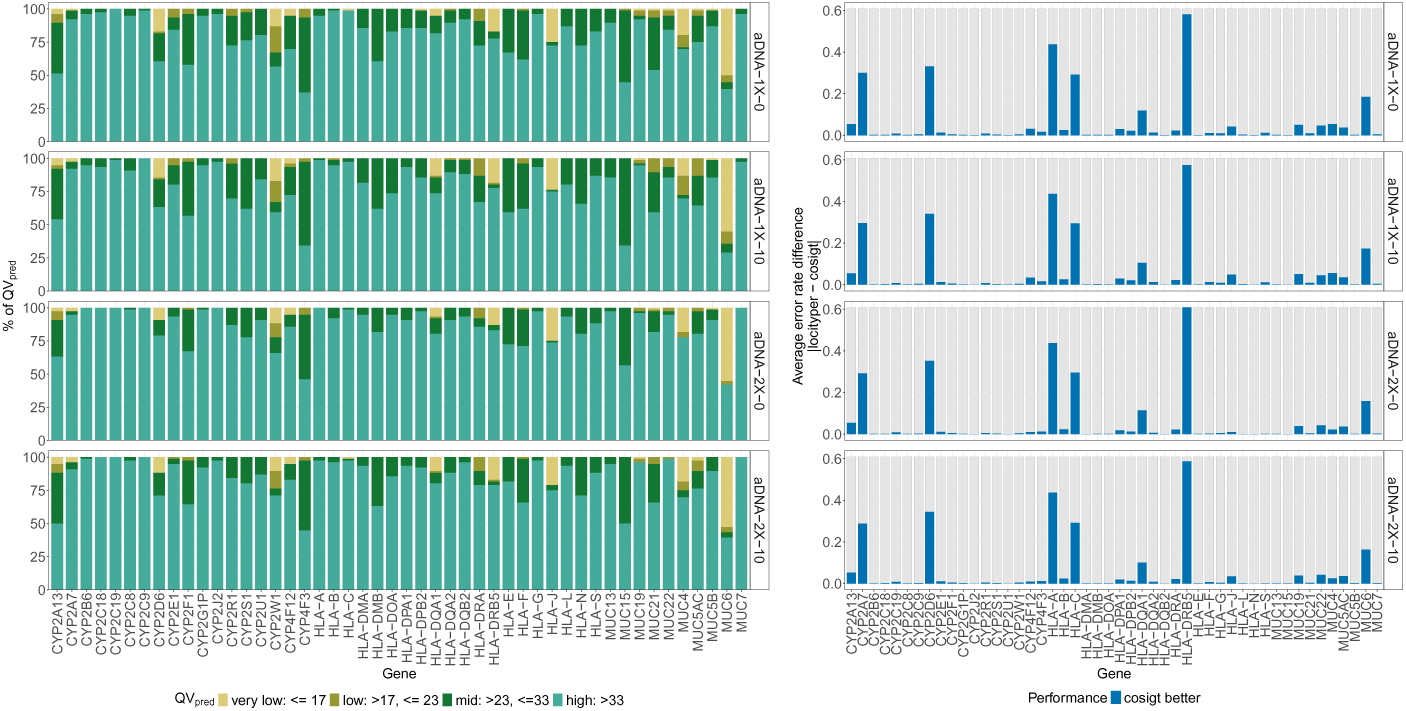
Gene-level COSIGT performance on simulated aDNA at variable coverage and contamination. Per-gene comparison of COSIGT and Locityper genotyping accuracy for 48 selected CMRGs on 38 simulated aDNA samples across four coverage and contamination conditions. Left: *QV*_pred_ distribution for COSIGT, showing the percentage of samples in each quality category (very low: *<*17, low: 17–23, mid: 23–33, high: *>*33; see Methods). Right: Average absolute error rate differences between COSIGT and Locityper. Rows represent different simulated datasets: aDNA at 1X coverage with 0% contamination (aDNA-1X-0), 1X with 10% contamination (aDNA-1X-10), 2X with 0% contamination (aDNA-2X-0), and 2X with 10% contamination (aDNA-2X-10). Genes are ordered alphabetically.

**Fig. S10:**
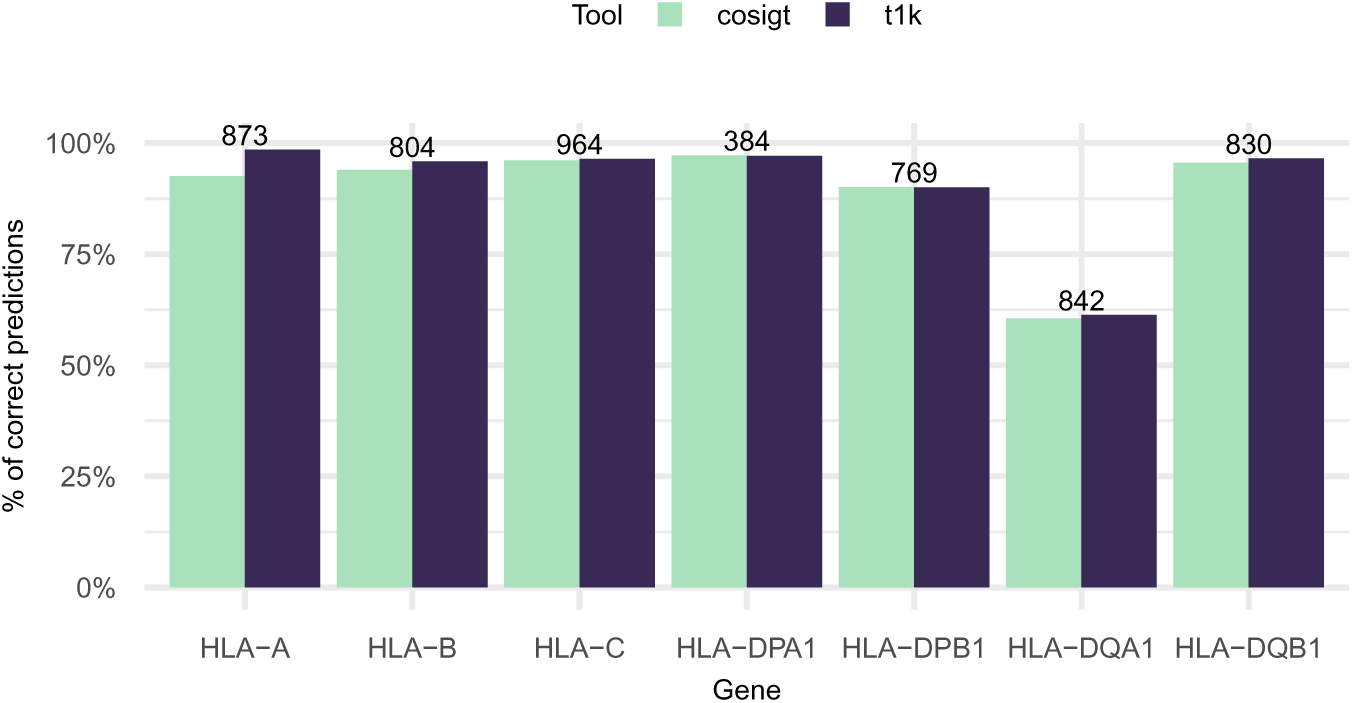
COSIGT performance on HLA typing. Haplotype-level concordance between short-read-based genotyping (COSIGT, T1K) and microarray-based imputation (HLA*IMP:02) for seven HLA genes (HLA-A, -B, -C, -DPA1, -DPB1, -DQA1, -DQB1) in a subset (*n* = 1,085) of the Moli-sani cohort, after filtering for alleles with quality less than or equal to 0 in T1K results. Numbers above bars indicate samples with predictions from all three methods and at least one reference haplotype present in the pangenome graph for the imputed genotype.

**Fig. S11:**
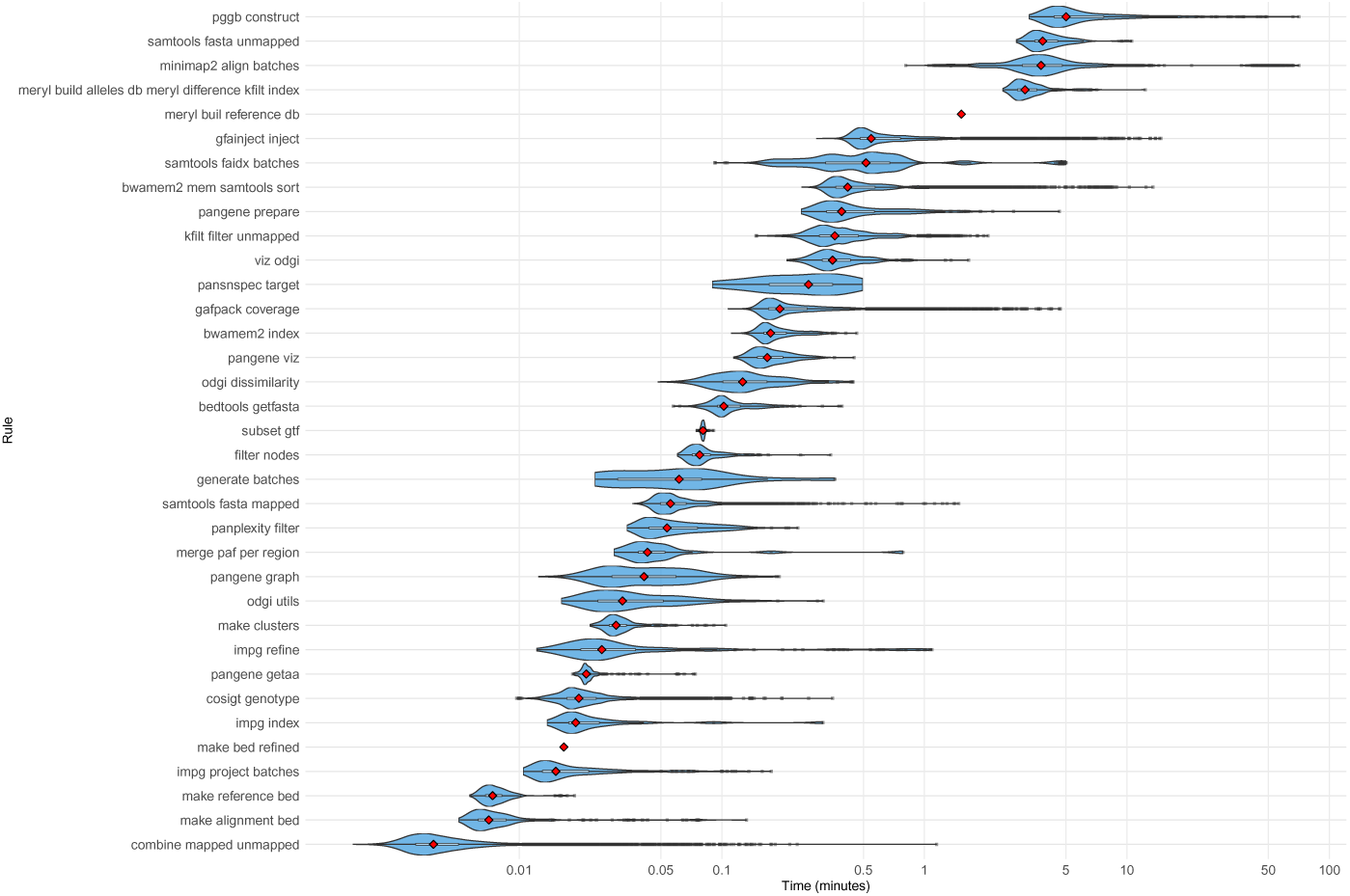
COSIGT runtime across pipeline steps. Violin plots show the distribution of the execution time (minutes) for each pipeline rule across jobs for the leave-zero-out analysis of SVs (see Methods). Boxplots (black) indicate quartiles, and red diamonds mark median values. X-axis uses a logarithmic scale. Rules are ordered by median execution time from slowest (top) to fastest (bottom).

**Fig. S12:**
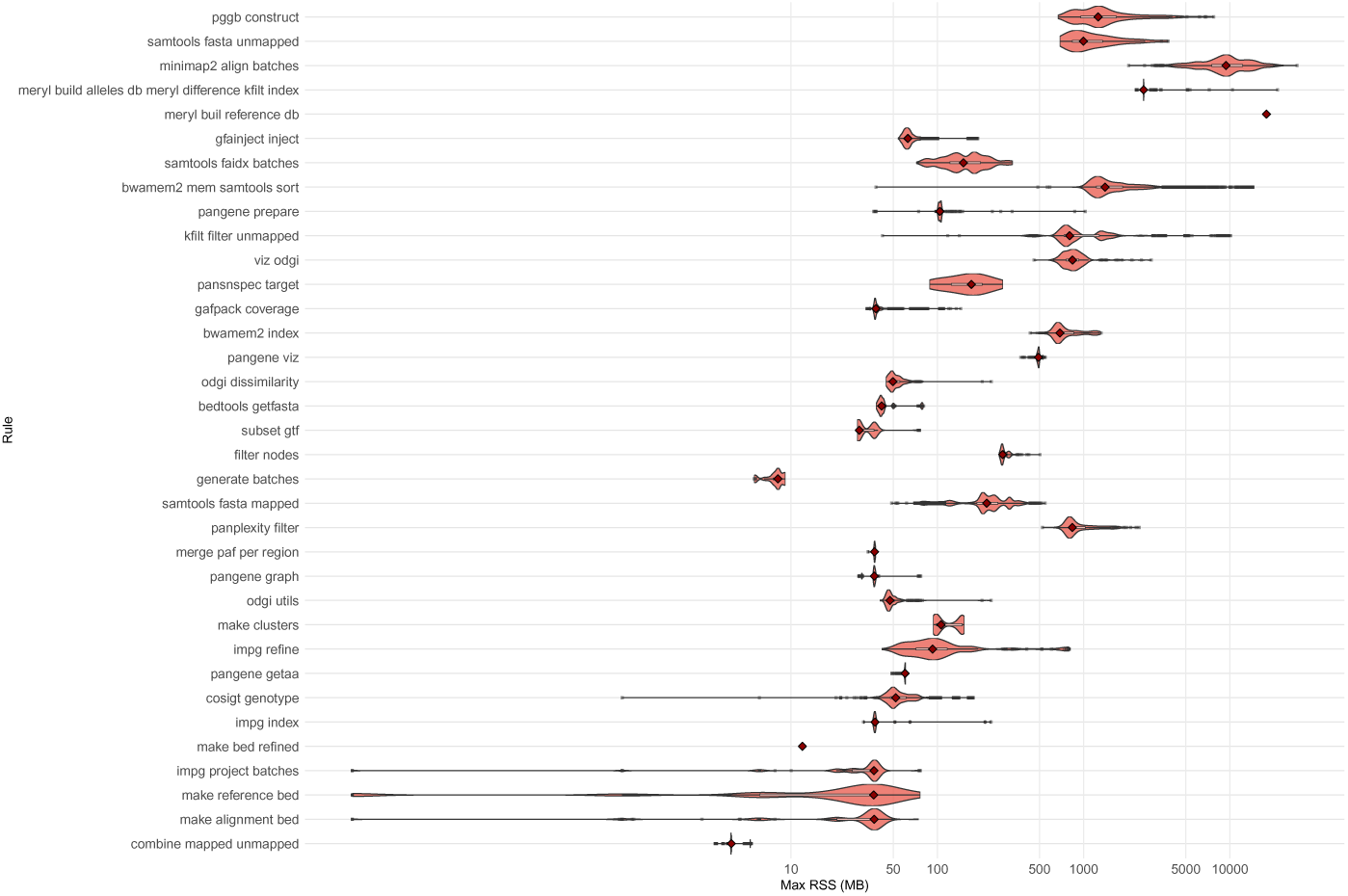
COSIGT memory usage across pipeline steps. Violin plots show the distribution of peak memory consumption (megabytes, MB) for each pipeline rule across jobs for the leave-zero-out analysis of SVs (see Methods). Boxplots (black) indicate quartiles, and red diamonds mark median values. X-axis uses a logarithmic scale. Rules are ordered by median execution time for consistency with Supplementary Figure S11.

### Supplementary C Supplementary Data Notes

#### C.1 Chromosome partitioning of haplotype-resolved assemblies

The COSIGT pipeline requires haplotype-resolved assemblies partitioned by chromosome to restrict computational analyses to relevant genomic regions. For this study, we used chromosome assignments provided by the HGSVCv3 and HPRCy1 consortia. However, such assignments may not always be available for newly generated assemblies or assemblies from alternative sources.

When reference-based chromosome annotations are unavailable, assemblies can be partitioned using sequence similarity to a reference genome. We provide a reference implementation in the COSIGT documentation (https://davidebolo1993.github.io/cosigtdoc/usecases/usecases.html) that uses Mash [46] distance to assign assembly contigs to their most likely chromosome of origin. Briefly, this approach: (1) computes k-mer sketches for both the reference genome and query assembly contigs using Mash, (2) calculates pairwise Mash distances between each contig and reference chromosomes, and (3) assigns each contig to the reference chromosome with the minimum distance.

For each contig, the best-matching reference chromosome is identified by selecting the alignment with the lowest Mash distance. Contigs assigned to the same chromosome are then concatenated into chromosome-specific FASTA files, maintaining the partitioned structure required by the pipeline. The complete workflow, including example commands and parameter settings, is available at https://davidebolo1993.github.io/cosigtdoc/usecases/usecases.html#option-2-starting-from-individual-haplotypes.

#### C.2 Incorporating alternative reference contigs for target regions

COSIGT accepts target regions as a standard BED file with genomic coordinates relative to the primary reference assembly. For basic analyses, a three-column BED format (chromosome, start, end) is sufficient. However, two optional columns can be added: column 4 for locus names and column 5 for comma-separated lists of alternative reference contigs corresponding to each target region.

Including alternative contigs is particularly important when working with reference genomes that contain alternate scaffolds, such as the GRCh38 reference used for 1000 Genomes Project alignments. These alternate sequences represent known variation (e.g., MHC haplotypes) and capture reads that would otherwise be unmapped or mismapped to the primary assembly. Failing to include these regions can result in incomplete read extraction and reduced genotyping accuracy, especially for highly polymorphic loci like the HLA genes.

To identify alternative contigs mapping to target regions, we provide a reference workflow in the COSIGT documentation (https://davidebolo1993.github.io/cosigtdoc/usecases/usecases.html). The approach: (1) extracts primary chromosomes and alternative scaffolds from the reference genome, (2) aligns alternative scaffolds to primary chromosomes using minimap2, (3) queries target regions against these alignments using impg to identify corresponding alternative contigs, and (4) merges primary and alternative coordinates into an extended BED file.

The resulting BED format enables COSIGT to extract reads mapped to both primary and alternative reference sequences, ensuring comprehensive coverage of sequence diversity at the target locus. For example, HLA typing analyses in this study used extended target regions that included both primary chromosome 6 coordinates and multiple HLA-specific alternative contigs from GRCh38, capturing reads that aligned to any representation of these loci. The complete workflow with detailed commands is available at https://davidebolo1993.github.io/cosigtdoc/usecases/usecases.html#incorporating-alternative-contigs.

## References

[1] Olson, N.D., Wagner, J., Dwarshuis, N., Miga, K.H., Sedlazeck, F.J., Salit, M., Zook, J.M.: Variant calling and benchmarking in an era of complete human genome sequences. Nature Reviews Genetics 24(7), 464–483 (2023). 10.1038/s41576-023-00590-0

[2] Liao, W.-W., Asri, M., Ebler, J., Doerr, D., Haukness, M., Hickey, G., Lu, S., Lucas, J.K., Monlong, J., Abel, H.J., Buonaiuto, S., Chang, X.H., Cheng, H., Chu, J., Colonna, V., Eizenga, J.M., Feng, X., Fischer, C., Fulton, R.S., Garg, S., Groza, C., Guarracino, A., Harvey, W.T., Heumos, S., Howe, K., Jain, M., Lu, T.-Y., Markello, C., Martin, F.J., Mitchell, M.W., Munson, K.M., Mwaniki, M.N., Novak, A.M., Olsen, H.E., Pesout, T., Porubsky, D., Prins, P., Sibbesen, J.A., Sirén, J., Tomlinson, C., Villani, F., Vollger, M.R., Antonacci-Fulton, L.L., Baid, G., Baker, C.A., Belyaeva, A., Billis, K., Carroll, A., Chang, P.-C., Cody, S., Cook, D.E., Cook-Deegan, R.M., Cornejo, O.E., Diekhans, M., Ebert, P., Fairley, S., Fedrigo, O., Felsenfeld, A.L., Formenti, G., Frankish, A., Gao, Y., Garrison, N.A., Giron, C.G., Green, R.E., Haggerty, L., Hoekzema, K., Hourlier, T., Ji, H.P., Kenny, E.E., Koenig, B.A., Kolesnikov, A., Korbel, J.O., Kordosky, J., Koren, S., Lee, H., Lewis, A.P., Magalhães, H., Marco-Sola, S., Marijon, P., McCartney, A., McDaniel, J., Mountcastle, J., Nattestad, M., Nurk, S., Olson, N.D., Popejoy, A.B., Puiu, D., Rautiainen, M., Regier, A.A., Rhie, A., Sacco, S., Sanders, A.D., Schneider, V.A., Schultz, B.I., Shafin, K., Smith, M.W., Sofia, H.J., Tayoun, A.N.A., Thibaud-Nissen, F., Tricomi, F.F., Wagner, J., Walenz, B., Wood, J.M.D., Zimin, A.V., Bourque, G., Chaisson, M.J.P., Flicek, P., Phillippy, A.M., Zook, J.M., Eichler, E.E., Haussler, D., Wang, T., Jarvis, E.D., Miga, K.H., Garrison, E., Marschall, T., Hall, I.M., Li, H., Paten, B.: A draft human pangenome reference. Nature 617(7960), 312–324 (2023). 10.1038/s41586-023-05896-x

[3] Asri, M., Chang, P.-C., Mier, J.C., Sirén, J., Eskandar, P., Kolesnikov, A., Cook, D.E., Brambrink, L., Hickey, G., Novak, A.M., Dorfman, L., Webster, D.R., Carroll, A., Paten, B., Shafin, K.: Pangenome-aware deepvariant (2025). 10.1101/2025.06.05.657102. Preprint

[4] Behera, S., Catreux, S., Rossi, M., Truong, S., Huang, Z., Ruehle, M., Visvanath, A., Parnaby, G., Roddey, C., Onuchic, V., Finocchio, A., Cameron, D.L., English, A., Mehtalia, S., Han, J., Mehio, R., Sedlazeck, F.J.: Comprehensive genome analysis and variant detection at scale using dragen. Nature Biotechnology 43(7), 1177–1191 (2024). 10.1038/s41587-024-02382-1

[5] Prodanov, T., Plender, E.G., Seebohm, G., Meuth, S.G., Eichler, E.E., Marschall, T.: Locityper enables targeted genotyping of complex polymorphic genes. Nature Genetics 57(11), 2901–2908 (2025). 10.1038/s41588-025-02362-4

[6] Rubinacci, S., Hofmeister, R.J., Sousa da Mota, B., Delaneau, O.: Imputation of low-coverage sequencing data from 150,119 uk biobank genomes. Nature Genetics 55(7), 1088–1090 (2023). 10.1038/s41588-023-01438-3

[7] Allentoft, M.E., Sikora, M., Refoyo-Martínez, A., Irving-Pease, E.K., Fischer, A., Barrie, W., Ingason, A., Stenderup, J., Sjögren, K.-G., Pearson, A., et al.: Population genomics of post-glacial western eurasia. Nature 625(7994), 301–311 (2024). 10.1038/s41586-023-06865-0

[8] Garrison, E., Guarracino, A., Heumos, S., Villani, F., Bao, Z., Tattini, L., Hagmann, J., Vorbrugg, S., Marco-Sola, S., Kubica, C., Ashbrook, D.G., Thorell, K., Rusholme-Pilcher, R.L., Liti, G., Rudbeck, E., Golicz, A.A., Nahnsen, S., Yang, Z., Mwaniki, M.N., Nobrega, F.L., Wu, Y., Chen, H., de Ligt, J., Sudmant, P.H., Huang, S., Weigel, D., Soranzo, N., Colonna, V., Williams, R.W., Prins, P.: Building pangenome graphs. Nature Methods 21(11), 2008–2012 (2024). 10.1038/s41592-024-02430-3

[9] The 1000 Genomes Project Consortium: A global reference for human genetic variation. Nature 526(7571), 68–74 (2015). 10.1038/nature15393

[10] Logsdon, G.A., Ebert, P., Audano, P.A., Loftus, M., Porubsky, D., Ebler, J., Yilmaz, F., Hallast, P., Prodanov, T., Yoo, D., Paisie, C.A., Harvey, W.T., Zhao, X., Martino, G.V., Henglin, M., Munson, K.M., Rabbani, K., Chin, C.-S., Gu, B., Ashraf, H., et al.: Complex genetic variation in nearly complete human genomes. Nature 644(8076), 430–441 (2025). 10.1038/s41586-025-09140-6

[11] Dilthey, A., Leslie, S., Moutsianas, L., Shen, J., Cox, C., Nelson, M.R., McVean, G.: Multi-population classical hla type imputation. PLoS Computational Biology 9(2), 1002877 (2013). 10.1371/journal.pcbi.1002877

[12] Song, L., Bai, G., Liu, X.S., Li, B., Li, H.: Efficient and accurate kir and hla genotyping with massively parallel sequencing data. Genome Research 33(6), 923–931 (2023). 10.1101/gr.277585.122

[13] Bolognini, D., Halgren, A., Lou, R.N., Raveane, A., Rocha, J.L., Guarracino, A., Soranzo, N., Chin, C.-S., Garrison, E., Sudmant, P.H.: Recurrent evolution and selection shape structural diversity at the amylase locus. Nature 634(8034), 617–625 (2024). 10.1038/s41586-024-07911-1

[14] Wick, R.R., Schultz, M.B., Zobel, J., Holt, K.E.: Bandage: interactive visualization of de novo genome assemblies. Bioinformatics 31(20), 3350– 3352 (2015). 10.1093/bioinformatics/btv383

[15] Garrison, E., Guarracino, A.: Unbiased pangenome graphs. Bioinformatics 39(1) (2022). 10.1093/bioinformatics/btac743

## Methods References

[16] Felix Mölder, Kim Philipp Jablonski, Brice Letcher, Michael B. Hall, Christopher H. Tomkins-Tinch, Vanessa Sochat, Jan Forster, Soohyun Lee, Sven O. Twardziok, Alexander Kanitz, Andreas Wilm, Manuel Holtgrewe, Sven Rahmann, Sven Nahnsen, and Johannes Köster. Sustainable data analysis with snakemake. F1000Research, 10:33, April 2021.

[17] Gregory M. Kurtzer, Vanessa Sochat, and Michael W. Bauer. Singularity: Scientific containers for mobility of compute. PLOS ONE, 12(5):e0177459, May 2017.

[18] conda contributors. conda: A system-level, binary package and environment manager running on all major operating systems and platforms.

[19] Heng Li. Minimap2: pairwise alignment for nucleotide sequences. Bioinformatics, 34(18):3094–3100, May 2018.

[20] Aaron R Quinlan and Ira M Hall. Bedtools: a flexible suite of utilities for comparing genomic features. Bioinformatics, 26(6):841–842, 2010.

[21] Andrea Guarracino, Simon Heumos, Sven Nahnsen, Pjotr Prins, and Erik Garrison. ODGI: understanding pangenome graphs. Bioinformatics, May 2022.

[22] Arang Rhie, Brian P Walenz, Sergey Koren, and Adam M Phillippy. Merqury: reference-free quality, completeness, and phasing assessment for genome assemblies. Genome Biology, 21(1):245, 2020.

[23] Petr Danecek, James K Bonfield, Jennifer Liddle, John Marshall, Valeriu Ohan, Martin O Pollard, Andrew Whitwham, Thomas Keane, Shane A McCarthy, Robert M Davies, and Heng Li. Twelve years of SAMtools and BCFtools. GigaScience, 10(2):giab008, 2021.

[24] Md Vasimuddin, Sanchit Misra, Heng Li, and Srinivas Aluru. Efficient architecture-aware acceleration of BWA-MEM for multicore systems. In 2019 IEEE International Parallel and Distributed Processing Symposium (IPDPS), pages 314–324. IEEE, 2019.

[25] Michael Hahsler, Matthew Piekenbrock, and Derek Doran. dbscan: Fast density-based clustering with R. Journal of Statistical Software, 91(1):1– 30, 2019.

[26] Adrien Oliva, Raymond Tobler, Bastien Llamas, and Yassine Souilmi. Additional evaluations show that specific BWA-aln settings still outper- form BWA-mem for ancient DNA data alignment. Ecology and Evolution, 11(24):18743–18748, December 2021.

[27] Heng Li. Protein-to-genome alignment with miniprot. Bioinformatics, 39(1), January 2023.

[28] Heng Li, Maximillian Marin, and Maha R Farhat. Exploring gene content with pangene graphs. Bioinformatics, 40(7), July 2024.

[29] David Porubsky, Xavi Guitart, DongAhn Yoo, Philip C. Dishuck, William T. Harvey, and Evan E. Eichler. SVbyEye: A visual tool to characterize structural variation among whole-genome assemblies. September 2024.

[30] Mikko Rautiainen, Sergey Nurk, Brian P Walenz, Glennis A Logsdon, David Porubsky, Arang Rhie, Evan E Eichler, Adam M Phillippy, and Sergey Koren. Telomere-to-telomere assembly of diploid chromosomes with Verkko. Nature Biotechnology, 41(10):1474–1482, 2023.

[31] Haoyu Cheng, Gregory T Concepcion, Xiaowen Feng, Haowen Zhang, and Heng Li. Haplotype-resolved de novo assembly using phased assembly graphs with hifiasm. Nature Methods, 18(2):170–175, 2021.

[32] Justin Wagner, Nathan D. Olson, Lindsay Harris, Jennifer McDaniel, Haoyu Cheng, Arkarachai Fungtammasan, Yih-Chii Hwang, Richa Gupta, Aaron M. Wenger, William J. Rowell, Ziad M. Khan, Jesse Farek, Yiming Zhu, Aishwarya Pisupati, Medhat Mahmoud, Chunlin Xiao, Byunggil Yoo, Sayed Mohammad Ebrahim Sahraeian, Danny E. Miller, David Jáspez, José M. Lorenzo-Salazar, Adrián Muñoz Barrera, Luis A. Rubio-Rodríguez, Carlos Flores, Giuseppe Narzisi, Uday Shanker Evani, Wayne E. Clarke, Joyce Lee, Christopher E. Mason, Stephen E. Lincoln, Karen H. Miga, Mark T. W. Ebbert, Alaina Shumate, Heng Li, Chen-Shan Chin, Justin M. Zook, and Fritz J. Sedlazeck. Curated variation benchmarks for challenging medically relevant autosomal genes. Nature Biotechnology, 40(5):672–680, February 2022.

[33] Martin Šošć and Mile Šikíc. Edlib: a C/C++ library for fast, exact sequence alignment using edit distance. Bioinformatics, 33(9):1394–1395, 2017.

[34] Owen D. Solberg, Steven J. Mack, Alex K. Lancaster, Richard M. Single, Yingssu Tsai, Alicia Sanchez-Mazas, and Glenys Thomson. Balancing selection and heterogeneity across the classical human leukocyte antigen loci: A meta-analytic review of 497 population studies. Human Immunology, 69(7):443–464, July 2008.

[35] Fernando A. Villanea, David Peede, Eli J. Kaufman, Valeria Añorve Garibay, Elizabeth T. Chevy, Viridiana Villa-Islas, Kelsey E. Witt, Roberta Zeloni, Davide Marnetto, Priya Moorjani, Flora Jay, Paul N. Valdmanis, María C. Ávila Arcos, and Emilia Huerta-Śanchez. The MUC19 gene: An evolutionary history of recurrent introgression and natural selection. Science, 389(6762), August 2025.

[36] M Ingelman-Sundberg. Genetic polymorphisms of cytochrome P450 2D6 (CYP2D6): clinical consequences, evolutionary aspects and functional diversity. The Pharmacogenomics Journal, 5(1):6–13, October 2004.

[37] Gabriel Renaud, Kristian Hanghøj, Eske Willeslev, and Ludovic Orlando. gargammel: a sequence simulator for ancient DNA. Bioinformatics, 33(4):577–579, 2017.

[38] Shifu Chen, Yanqing Zhou, Yaru Chen, and Jia Gu. fastp: an ultra-fast all-in-one FASTQ preprocessor. Bioinformatics, 34(17):i884–i890, 2018.

[39] Aaron McKenna, Matthew Hanna, Eric Banks, Andrey Sivachenko, Kristian Cibulskis, Andrew Kernytsky, Kiran Garimella, David Altshuler, Stacey Gabriel, Mark Daly, and Mark A DePristo. The Genome Analysis Toolkit: a MapReduce framework for analyzing next-generation DNA sequencing data. Genome Research, 20(9):1297–1303, 2010.

[40] Ryan Poplin, Pi-Chuan Chang, David Alexander, Scott Schwartz, Thomas Colthurst, Alexander Ku, Dan Newburger, Jojo Dijamco, Nam Nguyen, Pegah T Afshar, Sam S Gross, Lizzie Dorfman, Cory Y McLean, and Mark A DePristo. A universal snp and small-indel variant caller using deep neural networks. Nature Biotechnology, 36(10):983–987, September 2018.

[41] Kyle A. O’Connell, Zelaikha B. Yosufzai, Ross A. Campbell, Collin J. Lobb, Haley T. Engelken, Laura M. Gorrell, Thad B. Carlson, Josh J. Catana, Dina Mikdadi, Vivien R. Bonazzi, and Juergen A. Klenk. Accelerating genomic workflows using NVIDIA parabricks. BMC Bioinformatics, 24(1), May 2023.

[42] Brent S Pedersen and Aaron R Quinlan. Mosdepth: quick coverage calculation for genomes and exomes. Bioinformatics, 34(5):867–868, October 2017.

[43] Brent S. Pedersen, Preetida J. Bhetariya, Joe Brown, Stephanie N. Kravitz, Gabor Marth, Randy L. Jensen, Mary P. Bronner, Hunter R. Underhill, and Aaron R. Quinlan. Somalier: rapid relatedness estimation for cancer and germline studies using efficient genome sketches. Genome Medicine, 12(1), July 2020.

[44] Wenhan Lu, Laura D. Gauthier, Timothy Poterba, Edoardo Giacopuzzi, Julia K. Goodrich, Christine R. Stevens, Daniel King, Mark J. Daly, Benjamin M. Neale, and Konrad J. Karczewski. CHARR efficiently estimates contamination from DNA sequencing data. The American Journal of Human Genetics, 110(12):2068–2076, December 2023.

[45] Heng Li. A statistical framework for SNP calling, mutation discovery, association mapping and population genetical parameter estimation from sequencing data. Bioinformatics, 27(21):2987–2993, September 2011.

[46] Brian D. Ondov, Todd J. Treangen, Páll Melsted, Adam B. Mallonee, Nicholas H. Bergman, Sergey Koren, and Adam M. Phillippy. Mash: fast genome and metagenome distance estimation using minhash. Genome Biology, 17(1), June 2016.

